# Spectral Representation of EEG Data using Learned Graphs with Application to Motor Imagery Decoding

**DOI:** 10.1101/2022.08.13.503836

**Authors:** Maliheh Miri, Vahid Abootalebi, Hamid Saeedi-Sourck, Dimitri Van De Ville, Hamid Behjat

## Abstract

Electroencephalography (EEG) data entail a complex spatiotemporal structure that reflects ongoing organization of brain activity. Characterization of the spatial patterns is an indispensable step in numerous EEG processing pipelines within the setting of brain-computer interface systems as well as cognitive neuroscience. We present an approach for transforming EEG data into a spectral representation by using the harmonic basis of a graph structure that is learned from the data. The harmonic basis is obtained by integrating principles from graph learning and graph signal processing (GSP). First, we learn subject-specific graphs from each subject’s EEG data. Second, by eigendecomposition of the normalized Laplacian matrix of each subject’s graph, an orthonormal basis is obtained onto which each EEG map can be decomposed, providing a spectral representation of the data. We show that energy of the EEG maps is strongly associated with low frequency components of the learned basis, reflecting the smooth topography of EEG maps as expected. As a proof-of-concept for this alternative view of EEG data, we consider the task of decoding two-class motor imagery (MI) data. To this aim, the spectral representations are first mapped into a discriminative subspace for differentiating two-class data using a projection matrix obtained by the Fukunaga-Koontz transform (FKT), providing a minimal subspace from which features are extracted. An SVM classifier is then trained and tested on the resulting features to differentiate MI classes. The proposed method is evaluated on Dataset IVa of the BCI Competition III and its performance is compared to using features extracted from a subject-specific functional connectivity matrix and four state-of-the-art alternative methods. Experimental results indicate the superiority of the proposed method over alternative approaches, reflecting the added benefit of i) decomposing EEG data using data-driven, subject-specific harmonic bases, and ii) accounting for class-specific temporal variations in spectral profiles via the FKT. The proposed method and results underline the importance of integrating spatial and temporal characteristics of EEG signals in extracting features that can more powerfully differentiate MI classes.

## 1. Introduction

Electroencephalography (EEG) enables acquisition of brain activity at high temporal resolution, using multiple electrodes spanning the surface of the brain. The temporal and spatial structure of EEG data are both essential attributes to take into consideration when making interpretations and extracting features. Activity-induced electric fields get smeared as they transition from their source to the surface of the brain, and as such, the data has low spatial resolution and the channels are often highly correlated. Accounting for the underlying spatial organization in EEG data is therefore of marked importance. Graph signal processing (GSP) (Shuman et al., 2013; Ortega et al., 2018; Stanković et al., 2020), an emerging field that has attracted great interest across multiple disciplines, can be leveraged to account for the underlying structure in EEG data.

The fundamental idea in GSP is to apply signal processing procedures to data that reside on an irregular domain described by a graph, a construct consisting of a set of vertices and edges. GSP has in particular been adopted in a steadily increasing number of studies for characterization and processing of functional MRI data (Huang et al., 2018; Atasoy et al., 2017; Preti and Van De Ville, 2019; Abramian et al., 2021; Luppi et al., 2022; Behjat et al., 2015, 2022). For EEG data, a number of studies have shown promising results, namely, for dimensionality reduction (Tanaka et al., 2016; Kalantar et al., 2017), signal denoising (Cattai et al., 2021), and motor imagery (MI) decoding (Georgiadis et al., 2021; Cattai et al., 2022). Moreover, diffusion MRI-derived connectome harmonics have been used to characterize EEG data within a GSP setting, for tracking fast spatio-temporal cortical dynamics (Glomb et al., 2020b), their sparse representation (Rué-Queralt et al., 2021), and for source reconstruction (Rué-Queralt et al., 2022).

Despite the benefits of GSP, its successful application heavily relies on using a suitable graph that can represent an intrinsic underlying relation between the data elements. This is not available in many applications, such as for EEG data, in particular, if one is to consider solely an EEG dataset at hand; although graph vertices can be readily defined as being either recording electrodes (Pirondini et al., 2016; Georgiadis et al., 2021; Cattai et al., 2021) or regions from an atlas (Glomb et al., 2020b; Rué-Queralt et al., 2021), there exists no straight-forward definition of graph edges and edge weights. Given an ensemble set of data, graph learning (GL) techniques can be employed to infer a graph structure from data. Different GL methods have been proposed in the literature (Xia et al., 2021), a sub-category of which impose constraints on graph sparsity and signal smoothness (Dong et al., 2016; Kalofolias, 2016; Mateos et al., 2019). A number of recent neuroimaging studies have employed GL, namely, for subject identification (Gao et al., 2021), signal denoising (Einizade and Sardouie, 2022), and brain state identification (Ghoroghchian et al., 2020; Saboksayr et al., 2021).

Inspired by the promising results on the use of GSP and GL in brain mapping, here we propose a general scheme for GSP-based representing EEG signals on the spectra of graphs inferred via GL. In doing so, first, we learn a suitable graph from an ensemble set of EEG maps. Second, by employing GSP, we transform a desired set of EEG maps—which may not be necessarily the same as the set used to infer the graph—into a spectral representation, a space defined by the Laplacian harmonic basis of the learned graphs. This representation provides a compact and intuitive representation of EEG maps. A schematic overview of the proposed method is illustrated in Figure 1.

**Figure 1:**
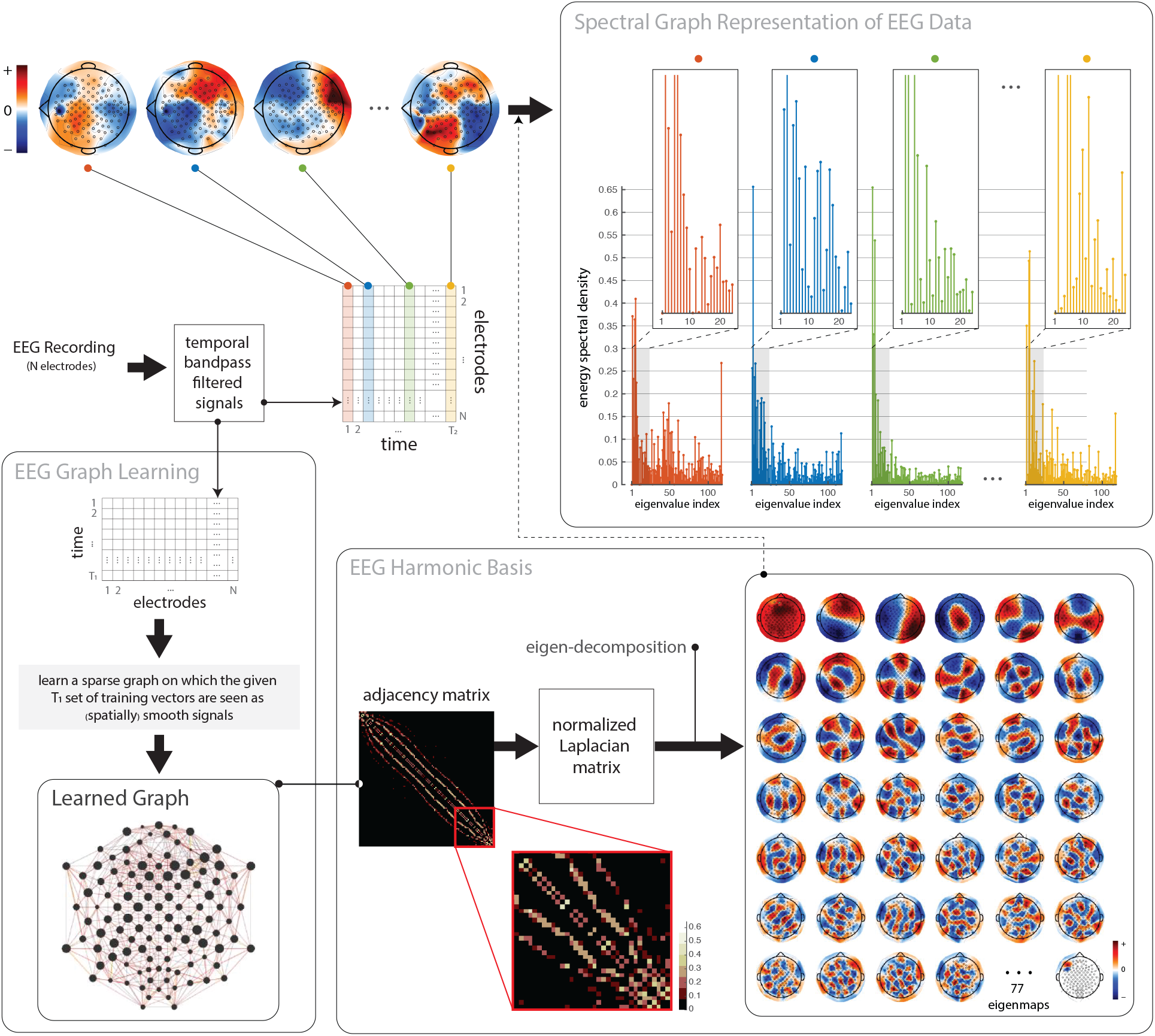
A schematic overview of the proposed methodology for spectral representation of EEG data on the harmonic basis of learned graphs. A *T*_1_ set of *N*electrode EEG maps are used to learn a graph, using which an EEG harmonic basis is derived. A *T*_2_ set of *N*-electrode EEG signals can then be decomposed using the learned harmonic basis; decomposition of four representative signals (time points) from a set are shown. The *T*_1_ and *T*_2_ set of measurements may be identical or may differ, depending on the application at hand.

To showcase the applicability of the proposed EEG mapping, we consider the problem of MI task decoding (Pfurtscheller and Neuper, 2001). Discrimination of mental states from EEG data is a challenging task, for which numerous methods have been proposed (Singh et al., 2021; Mirjalili et al., 2022). One class of methods aims at extracting features from the temporal evolution of the data, in time or frequency domain (Abootalebi et al., 2009; Kim et al., 2018; Lee et al., 2019; Gonzalez-Astudillo et al., 2021; Gwon and Ahn, 2021), whereas an alternative class aims at extracting spatial features from EEG maps (Ramoser et al., 2000; Lotte and Guan, 2010; Wang et al., 2020; Cherloo et al., 2021; Hou et al., 2022). Adaptive classifiers, matrix and tensor classifiers, transfer learning, and deep learning are among other methods that have been more recently proposed (Lotte et al., 2018; Peterson et al., 2021; Dose et al., 2018). Here we propose a method that is comprised of four stages. First, we use graph learning to infer subject-specific EEG graphs. Second, by interpreting EEG maps as time-dependent graph signals on the learned graphs, we transform the data into a spectral representation. Third, we derive a discriminative spectral graph subspace that specifically aims at differentiating two-class data, using a projection matrix obtained by the Fukunaga-Koontz transform (FKT) (Fukunaga, 2013); the transform takes in to account differences in temporal covariation of spectral profiles in the two classes, in the same spirit as methods that function on spatial features of the electrode space (Lotte and Guan, 2010; Wang et al., 2020; Georgiadis et al., 2021; Cohen, 2022). Fourth, we treat the variance of representations within the subspace as features, which is in turn used to train and test a binary classifier.

The remainder of this paper is structured as follows. In Section 2, we provide an overview of fundamental concepts from GSP, GL, and FKT, tailored for the EEG-based application at hand. In Section 3, we first study the learned graphs and their associated harmonic bases, highlighting differences across subjects as well with respect to graphs obtained via correlation-based functional connectivity. We then show that energy of EEG maps is strongly associated with low frequency components of the learned bases, providing a sparse representation of the data. Finally, we present MI-task decoding results using the proposed method, also bench marking against alternative state-of-the-art methods. In Section 4, we present a discussion around the results, highlight limitations, and provide an outlook into potential future work.

## 2. Materials and Method

### 2.1. Dataset

To evaluate the proposed method, EEG data from the publicly available BCI Competition III-Dataset IVa (Blankertz et al., 2006) were used. The data, comprising of two classes of motor imagery EEG signals, were recorded from five healthy subjects (labeled as aa, al, av, aw, and ay) using 118 electrodes that were installed with the electrode arrangement in the extended international 10/20-system at a sampling rate of 100 Hz. A total of 280 visual cues of length 3.5 seconds were presented to subjects, interleaved with rest interval of random lengths 1.75 to 2.25 seconds. Despite the limited number of subjects, the dataset is very rich in that it includes a large number of trials per subject. This makes the dataset very suitable for use within a machine learning setting, and as a benchmark in many studies. During the presentation of target cues, subjects were asked to perform right hand or right foot motor imageries. 140 trials were acquired for each class. According to the competition instructions the trials were divided into training and test sets in each class. The set sizes differed across the five subjects. For the first two subjects most trials are labeled (60% and 80%, respectively), whereas for the other three subjects 30%, 20%, and 10% of the trials are labeled, respectively; the remaining trials are used as test sets^1^. Given the difference in the size of the training and test sets across subjects, classification is more challenging on subjects av, *aw*, and *ay*, which have smaller training sets.

### 2.2. Graph Signal Processing Fundamentals

Let 𝒢 = (**𝒱, ℰ, A**) denote a weighted, undirected graph, where **𝒱** = {1, 2, …, *N*} represents the graph’s finite set of *N* vertices (nodes), **ℰ** denotes the graph’s edge set, i.e., pairs (*i, j*) where *i, j* ∈ ***𝒱***, and **A** is a symmetric matrix (*A*_*i, j*_ = *A*_*j,i*_) that denotes the graph’s weighted adjacency matrix. The weights in the adjacency matrix indicate the strength of the connection, or similarity between two corresponding vertices, therefore, *A*_*i, j*_ = 0 if there is no connection/similarity between vertices *i* and *j*. It is assumed that there are no self-loops in the graph, i.e., *A*_*i,i*_ = 0. The graph’s combinatorial Laplacian matrix is defined as

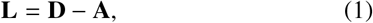

where **D** is the diagonal matrix of vertex degrees with its elements given as *D*_*i,i*_ = ∑_*j*_ *A*_*i, j*_, and the graph’s symmetrically normalized Laplacian matrix is defined as:

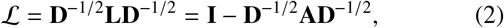

where **I** is the identity matrix. Since ℒ is a real, symmetric, and positive semi-definite, it can be diagonalized via its eigenvalue decomposition as:

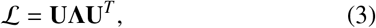

where *T* denotes the transpose operator, **U** = [**u**_1_, **u**_2_, …, **u**_*N*_] is an orthonormal matrix concatenating the eigenvectors **u**_*k*_ ∈ ℝ^*N*^ in its columns, and **Λ** is a diagonal matrix that stores the corresponding real, and non-negative eigenvalues 0 = *λ*_1_ ≤*λ*_2_≤ … ≤*λ*_*N*_≤ 2. The eigenvalues define the graph Laplacian spectrum, and the corresponding eigenvectors form an orthonormal harmonic basis (Chung, 1997). In the following, we interchangeably use the terms *eigenvector, harmonic*, and *eigenmap*, all referring to the eigenvectors of the normalized graph Laplacian matrix; in particular, we refer to eigenmaps when illustrating EEG harmonics as spatial maps. Graph Laplacian eigenvectors associated to larger eigenvalues entail a larger extent of variability. Specifically, eigenvalues of the graph Laplacian matrix can be seen as an extension of frequency elements that define the Fourier domain in classical signal processing (Ortega et al., 2018).

Let **f** ∈ ℝ^*N*^ denote a graph signal, that is, a real signal defined on the vertices of 𝒢 whose *n*-th component represents the signal value at the *n*-th vertex of 𝒢. By using the Laplacian eigenvectors, **f** can be transformed into its spectral representation, commonly referred to as the graph Fourier transform (GFT) of **f**, denoted 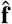, as:

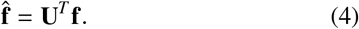

The GFT satisfies Parseval’s energy conservation relation, i.e., 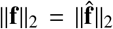 (Behjat et al., 2016), indicating that the energy of the signal can be computed equally in either the vertex domain or the spectral domain of 𝒢. Given the orthonormality of the Laplacian eigenvectors, the inverse GFT of 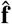 is obtained as:

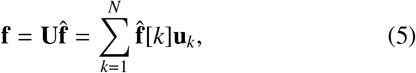

showing that by synthesizing **f** as a weighted sum of orthogonal graph frequency components **u**_*k*_, 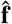 entails the degree of spatial variability of **f** over the 𝒢.

To better understand the notion of frequency on graphs, the total variation (TV) of a graph signal **f** on graph 𝒢 can be quantified using **L** as (Von Luxburg, 2007):

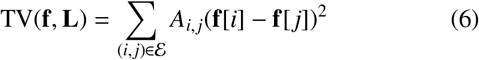

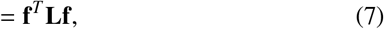

where the proof of the second equality is given in Appendix 1. Larger values of TV(**f, L**) indicate greater changes of **f** on 𝒢, i.e., higher spatial variability, and thus, lower spatial smoothness; this notion of smoothness is leveraged in Section 2.3 for learning graphs. Alternatively, instead of using **L**, TV can be computed in a similar way as in (7) using ℒ, which, in particular, provides an intuitive interpretation in the spectral domain as:

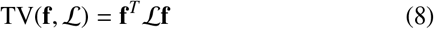

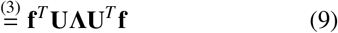

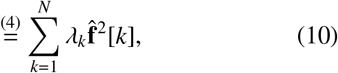

showing that signals that are spatially smooth on the graph— i.e., have the majority of their energy in the lower end of the spectrum—have a low TV whereas signals that exhibit higher order spatial fluctuations entail a lager TV; an alternative formulation of TV(**f**, ℒ) similar to (6) is given in Appendix 2. Moreover, by viewing each of the eigenvectors **u**_*k*_ as a graph signal, we have:

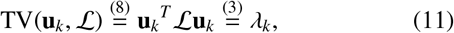

showing that each eigenvalue is a quantification of the extent of variability of its corresponding eigenvector. Alternatively, a more simplistic, and intuitive quantification of the extent of variability of eigenvectors is given by computing a weighted zero crossings (WZC) measure (Petrantonakis, 2021) as:

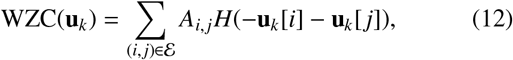

where *H*(·) is the Heaviside step function.

### 2.3. Learning Graphs from Smooth Signals

Effective use of various GSP algorithms relies on using suitable graphs that can capture subtle intrinsic organizational features within graph signals. However, in many applications, a definition of a graph is not readily available. Graph learning (GL) techniques can be used to estimate a graph structure from an available set of data. A class of GL methods enforce data smoothness—that is, graph signals should be smooth on the learned graph (Dong et al., 2016; Kalofolias, 2016). A graph signal is smooth on a given graph if graph signal elements that are connected via an edge with high weight exhibit small differences, whereas larger differences are observed between elements that are either not connected via an edge or connected via an edge with a small weight.

Let **F** = [**f**_1_, **f**_2_, …, **f**_*M*_] ∈ ℝ ^*N*×*M*^ denote a matrix that stores a set of *M* measurements on a domain with *N* elements, and let **Z** denote an *N*× *N* matrix with entries that represent the Euclidean distance between pairs of rows in **F**, i.e., *Z*_*i, j*_ = ll **F**_*i*,:_ −**F** _*j*,:_ ll_2_, where **F**_*k*,:_ denotes the *k*-th row of **F**, that is measurements from the *k*-th element. The objective is to derive an organizational relation between the rows of **F**, in the form of a graph. Given a graph 𝒢 with *N* vertices, the overall smoothness of the columns of **F**—i.e., graph signals on 𝒢—can be computed using (7) as:

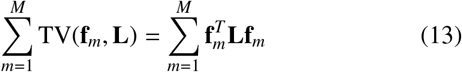

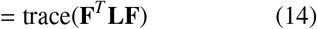

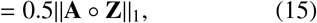

where ° denotes the Hadamard product, and **A** denotes the graph’s adjacency matrix; the proof of the last equality is given in Appendix 3. The smaller the value given by (15), the smoother is **F** on 𝒢; that is, for **F** to be smooth on graph 𝒢, non-zero elements in **A** should maximally match small-value elements in **Z**, meaning that signal values at vertices connected with an edge should minimally differ as quantified by the distance measure. Using this measure of smoothness, a graph can be learned via solving (Kalofolias, 2016):

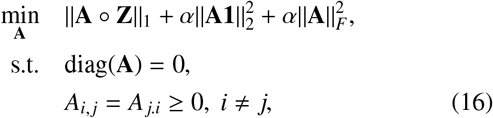

where *α* is regularization parameter, || · ||_*F*_ denotes the Frobenius norm and **1** = [1, …, 1]^*T*^. The terms 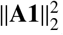 and 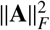 control sparsity by minimizing vertex degrees and shrinking edge weights, respectively, whereas the imposed constraints ensure finding a valid graph adjacency matrix. Alternatively, the objective function in (16) can be improved by replacing the *ℓ*_2_-norm with a logarithmic barrier on the vertices degree vector as:

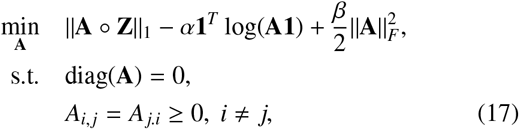

where the second term ensures graph degrees to be positive, thus improving the overall connectivity of the graph, and moreover, ensures each vertex to have at least one edge. *α* and *β* are regularization parameters; intuitively, smaller values of *β* yield sparser graphs by penalizing edges between vertices with larger *Z*_*i, j*_ (Kalofolias, 2016). In the following, we refer to the graph learning approaches given in (16) and (17) as the ℓ_2_-penalized and log-penalized methods, respectively.

### 2.4. Two-Class Discriminative Subspace via Simultaneous Diagonalization of Covariance Matrices

After defining a brain graph, the graph spectral representations of the EEG signals were considered to find a discriminative subspace for two-class (right hand and right foot) MI classification. To this end, inspired by methods presented in (Itani and Thanou, 2021; Subbaraju et al., 2017), we use the FKT (Fukunaga and Koontz, 1970; Fukunaga, 2013), which is based on simultaneous diagonalization of two covariance matrices (Cohen, 2022). For graph signal **f** defined on 𝒢, let 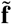 denote the de-meaned and normalized version of **f** obtained as (Behjat et al., 2021):

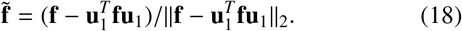

Given two signal classes, *j* = 1, 2, let 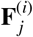 denote an *N* × *M* matrix of the EEG data for trial *i* of class *j*, wherein the element on the *n*-th row and *m*-th column corresponds to the signal value at electrode/vertex *n* at time point *m*; i.e., each column in 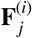 represents a graph signal. Let 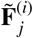 denote the de-meaned and normalized version of 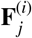, wherein each column is normalized as in (18), and let 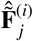 denote the GFT of 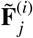, where the GFT is applied on each column of 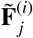. By leveraging the FKT, the goal is to map 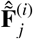 to a subspace in which the class type, i.e., *j*, can be inferred. This is done by considering the covariance structure of the GFT coefficients. Let

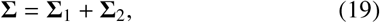

where ∑ _*j*_ denotes the across-trial ensemble averaged covariance matrix of class *j*, defined as:

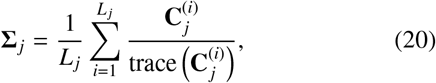

where *L*_*j*_ denotes the number of trials in class *j*, and 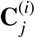 is the covariance matrix of the GFT coefficients of trial *i* of class *j*, obtained as:

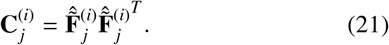

As **∑** is positive definite, it can be eigendecomposed as

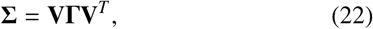

where **V** is the matrix of eigenvectors and Γ is the diagonal matrix of the corresponding eigenvalues, using which a whitening transform **P** is obtained as:

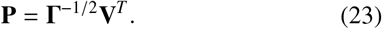

By whitening **∑**, the variances in the space spanned by **V** will become equal, i.e.:

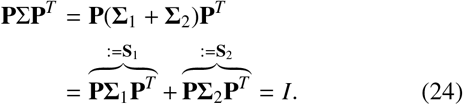

Consequently, eigenvalue decomposition of **S**_1_ and **S**_2_ gives:

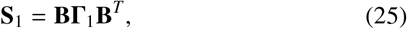

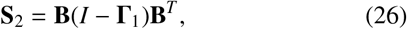

where **B** denotes the eigenvectors, which are the same for both **S**_1_ and **S**_2_, and their corresponding eigenvalues are complementary; i.e., by sorting the eigenvalues in descending order, the eigenvector associated with the largest eigenvalue of **S**_1_ is associated with the smallest eigenvalue of **S**_2_. Therefore, the combination of a small subset of the first and last eigenvectors of **B** provides a discriminatory transform for differentiating the two classes. Finally, the overall projection matrix can be obtained as:

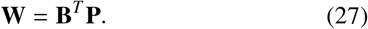

Using **W** = [**w**_1_, **w**_2_, …, **w**_*N*_]^*T*^, the GFT coefficients of a given EEG trial, i.e, 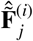, can be transformed to a feature vector **y** ∈ ℝ^*P*^ as:

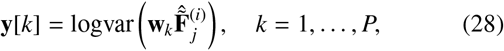

where *P* =| 𝒫| with 𝒫 ⊂ {1, 2, …, *N*} being a subset of the first and last GFT indices. Results presented in this work are for 𝒫 { = 1, *N*}, thus, resulting in a two-dimensional discriminative GFT subspace, wherein the variance of the first feature is maximized in the first class while being minimized in the second class and vice versa. For the interested reader, a mathematical analysis of using EEG maps as inputs to FKT, and the relation between resulting FKT filters to those presented here is given in Appendix 4.

### 2.5. Proposed Method for MI Task Decoding

The proposed method for EEG-based MI task decoding is illustrated as a block diagram in Figure 2. The training and test EEG signal sets for each subject are initially preprocessed, and then fed into the training and test phases, respectively. As temporal preprocessing, for each trial, we used the time points within the 0.5-2.5 second interval after the visual cue to construct graph signals; this 2-second interval has been previously used in related works (Wang et al., 2020; Cherloo et al., 2021; Georgiadis et al., 2021). Motor activity, be it real or imagined, modulates the sensorimotor mu rhythm (8-13 Hz) and beta rhythm (13-30 Hz), therefore, we filtered the extracted signal with a third-order Butterworth filter with a passband of 8-30 Hz. Graph signals were then extracted from these filtered signals; in particular, we defined one graph signal per time instance, i.e., each signal represents EEG values across the 118 electrodes, which, thus, resulted in *M* = 200 graph signals per trial.

**Figure 2:**
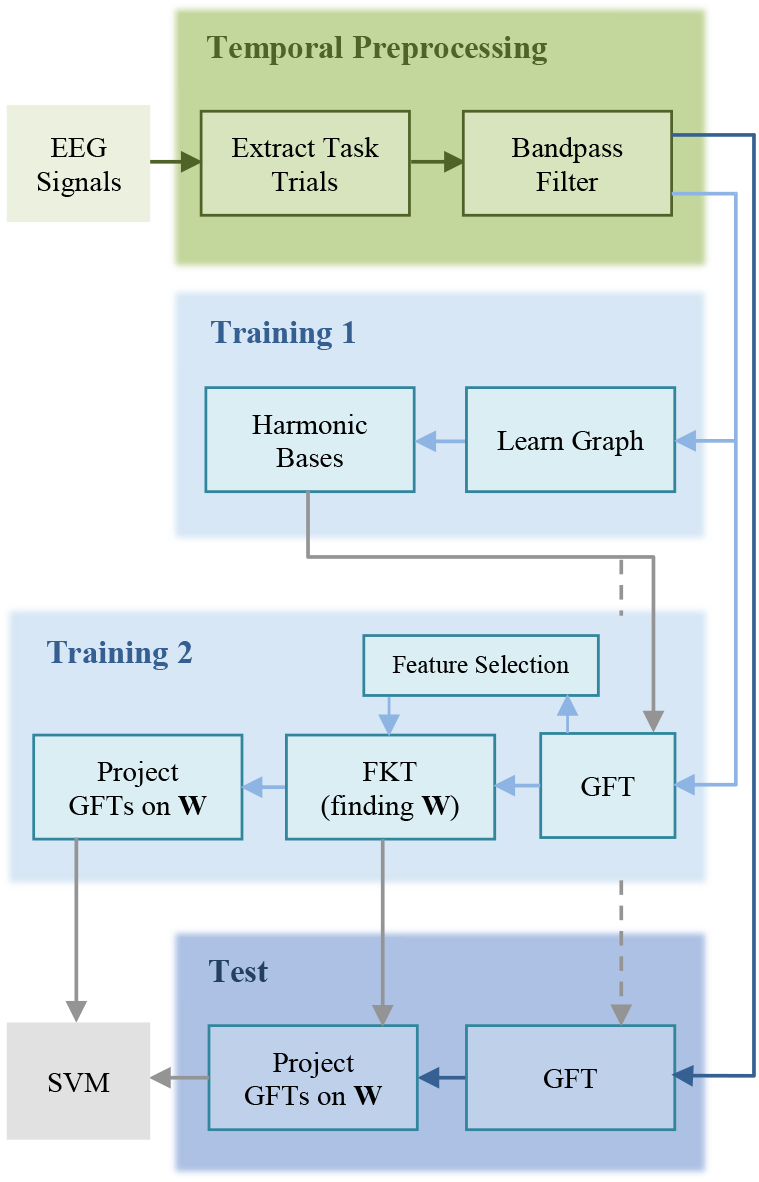
Block diagram of the proposed method for MI task decoding.

#### 2.5.1. EEG as Graph Signals

In the training phase, we modelled the structure of the brain of each subject as a graph, in which vertices corresponded to the EEG electrodes and edges were defined by estimating the graph’s weighted adjacency matrix using the log-penalized and *ℓ*_2_-penalized graph learning frameworks. As a means of comparison, we also defined a fully connected correlation graph in which edge weights were defined based on the degree of functional connectivity between electrode pairs; that is, for each electrode pair, the absolute value of the Pearson correlation coefficient between their time courses was defined as the edge weight, reflecting an estimate of the overall statistical dependency between the two electrodes (Miri et al., 2022). The large number of recording electrodes in the studied EEG dataset makes the dataset a good choice for graph-based analyses, providing the means to study the spatial organization and interaction of different cortical areas while also considering the fast temporal dynamics. In order to investigate the role of the brain structure (the physical distance between the vertices), the performance of two distance-based graphs was also investigated, in which, the Euclidean distance between the electrodes was utilized as the weight of the graph edges. The results of using these distance-based graphs are provided in the supplementary material for comparison.

#### 2.5.2. EEG Feature Extraction via GSP and FKT

For each graph, the eigenvectors of ℒ were used to compute the GFT of each graph signal. Using FKT, a transformation matrix **W** that maps the GFT coefficients into a discriminative graph spectral subspace was then derived. The mapped data were then treated as discriminative features. To determine the most effective graph frequency harmonics for classifying the EEG signals, a feature selection algorithm was used; we ranked the GFT coefficients based on their energy using the Wilcoxon statistical test. GFT Coefficients with higher ranks correspond to more distinctive features. The number of selected features for each subject was determined using 10-fold cross-validation.

#### 2.5.3. MI Classification and Evaluation

The classifier was trained using labelled training data, where labels indicate the class of each trial, and classification performance was evaluated with the labelled test data. The projection matrix **W** and the index of GFT coefficients that provided the most discriminative features were derived in the training phase and consequently used in the test phase. The logarithm of variance of the projected GFT coefficients on **W** were used as features to train a support vector machine (SVM) classifier with a linear kernel. Since this projection maximizes the variance of the signals from one class while minimizing it for the signals from the other class, it provides discriminative features for classification. We used SVM due to its overall superior robustness and efficiency in the BCI applications compared to other classifiers (Lotte et al., 2018). The linear kernel was selected for its simplicity and low computational cost.

## 3. Results

Figure 3(a) shows the arrangement of the 118 electrodes on the head; a map showing the correspondence between electrode positions and anatomical regions is provided in supplementary material Figure S1. Figure 3(b) shows the adjacency matrices of the three graphs for subject aa. Both the learned graphs are significantly sparser than the correlation graph, a result of sparsity-inducing terms used in the learning process. In contrast, the correlation graph is a complete graph, with edge weights that are substantially larger than those of the two learned graphs. Figure 3(c) presents schematic views of the three different graphs across the five subjects. The graphs of three subjects—aa, al, and aw—manifest approximately similar patterns, wherein a large number of edges are concentrated in the frontal and parietal lobes. In the other two subjects—av, and ay—edges are more broadly distributed across the brain. The number of graph edges are comparable between the learned graphs but are differently scaled for the correlation graph due to the large difference between the degree distributions. The correlation graph is a complete graph as it is defined based on the correlation of all electrode pairs, whereas the two learned graphs are notably sparse. A quantitative comparison of the graphs is presented in Table 1. It shows the connection density in the studied graphs which is the proportion of the number of graph edges relative to the total possible number of connections that could be formed in the graph. The graph learning methods preserve the connectivity of the graphs by using a lower number of edges compared to the fully connected correlation graphs. Moreover, the log-penalized method yields sparser learned graphs compared to the *ℓ*_2_-penalized method. The sparsity patterns of the learned graphs are compared in Figure 3(d). In all five subjects, the edges of the log-penalized graph are a subset of the edges of their corresponding *ℓ*_2_-penalized graph, but nevertheless, edge weights notably differ between the two designs. The differences between the weights of the common edges were obtained by subtracting the *ℓ*_2_-penalized edge weights from the log-penalized ones, results shown in Figure 3(e). The differences are positive values in all subjects, reflecting higher edge weights in log-penalized graphs. The highest edge weights are for subject av and the lowest values are for subject aw which also have the highest and the lowest connection densities in the learned graphs, respectively (cf. Table 1). For comparison, distance-based graphs as well as their adjacency matrices are shown in Figure S2 in the supplementary material. The distribution of nodal degrees, edge weights, and nodal strengths are shown in Figure 4. Nodal degrees of the correlation graphs are larger than that of the learned graphs, which is because i) they are complete graphs (cf. Table 1), and ii) the edge weights are in general greater than those of the learned graphs (cf. Figure 3(b)), which can also be visually inferred from Figure 3(c).

**Table 1:** Connection density of the three studied graph types for each subject and on average across all subjects.

| Graphs | aa | al | av | aw | ay | Mean±std |
| --- | --- | --- | --- | --- | --- | --- |
| log-penalized | 0.12 | 0.12 | 0.16 | 0.09 | 0.1 | 0.12 ± 0.03 |
| $\ell_2$-penalized | 0.18 | 0.19 | 0.22 | 0.16 | 0.16 | 0.18 ± 0.03 |
| correlation | 1 | 1 | 1 | 1 | 1 | 1 |

**Figure 3:**
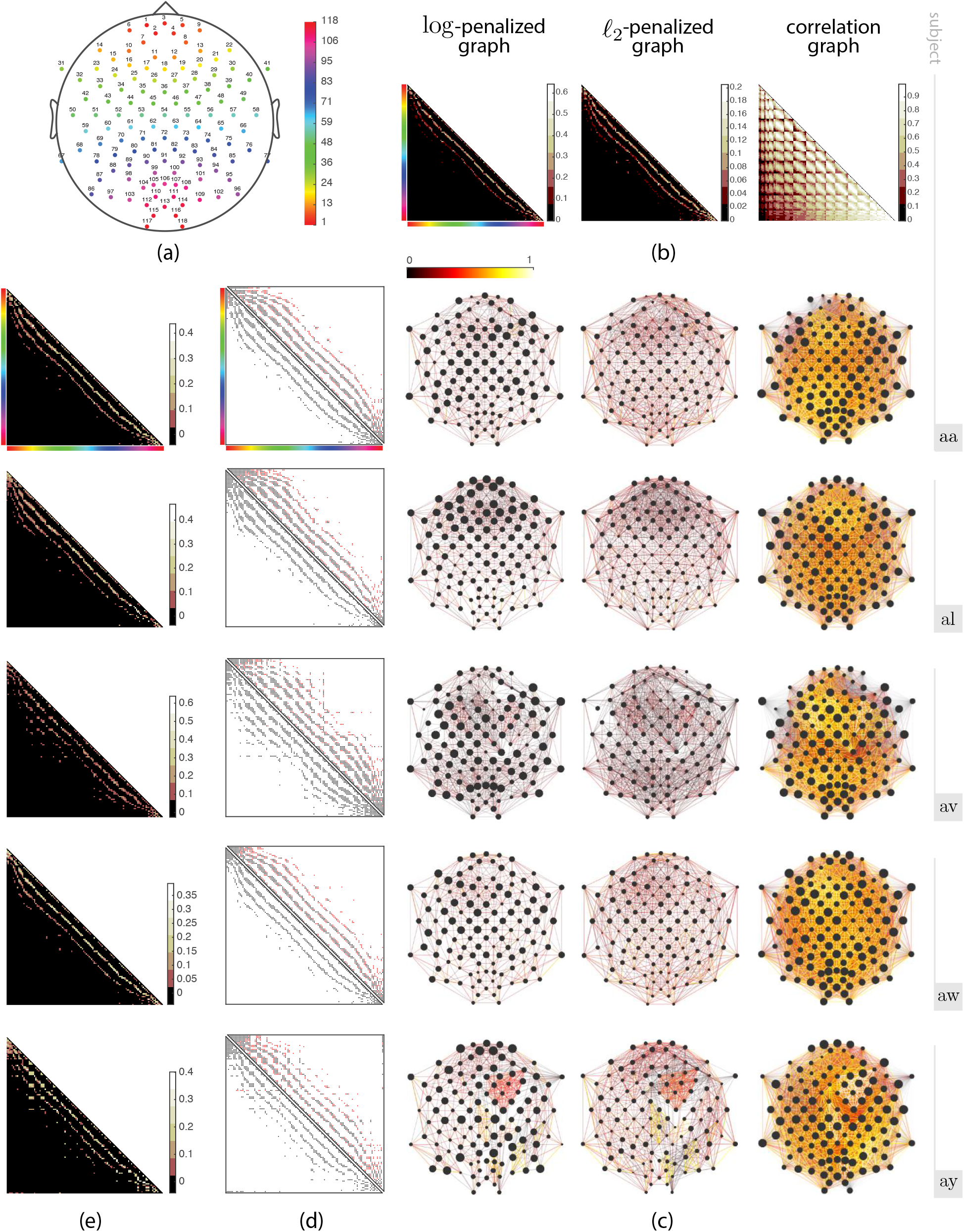
(a) Arrangement of EEG electrodes and their associated graph vertex indices. For better interpretation of graph matrices, a colormap is added to enable linking the row/column order of the matrices to the electrode layout; in subplots (b), (d) and (e), a representative matrix is appended with the colormap. (b) Adjacency matrices of the log-penalized, *ℓ*_2_-penalized, and correlation graphs for subject aa. (c) Schematic representation of the three different graphs, across five subjects. Edge widths and colors reflect edge weights, and vertex sizes reflect nodal degrees. For better visualization, only the top 50% of edges that have the largest weights are shown in the correlation graph. (d) Comparison of the sparsity pattern of the log-penalized graph (lower triangular segment) and the *ℓ*_2_-penalized graph (upper triangle segment) learned for each subject; edges that are common between the two graphs are shown as black dots, whereas edges that are unique to each design are shown as red dots. (e) Difference between edge weights of edges that are common between the log-penalized and *ℓ*_2_-penalized graph of each subject.

**Figure 4:**
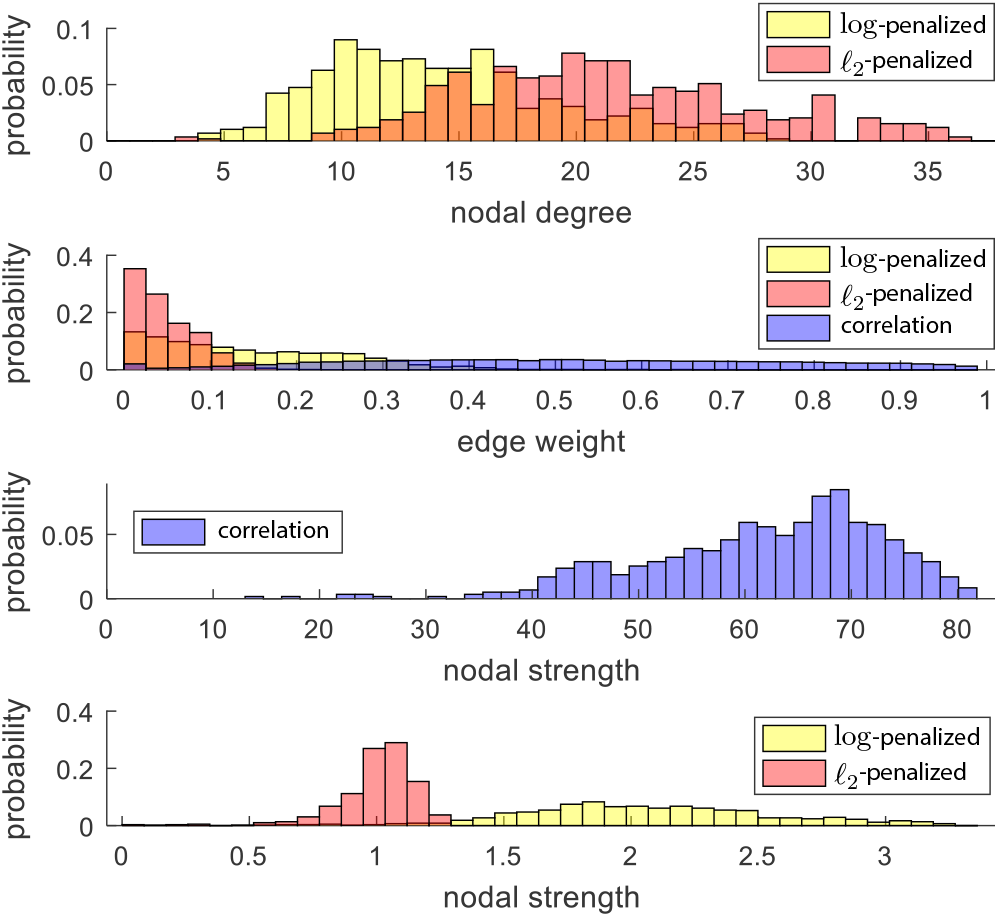
Distribution of nodal degrees, edge weights, and nodal strengths of the three studied graphs; for each graph type, for each of the three measures, values were pooled together across subjects and a probably distribution was then computed.

Distribution and histogram of the normalized Laplacian eigenvalues for three graphs of subject aa are shown in Figure 5(a). Most of the eigenvalues in the correlation graph are concentrated around the value of one, whereas the eigenvalues of the learned graphs, especially the log-penalized graph, gradually increase, and are more widely distributed along the spectrum. Figure S3 in the supplementary material shows the normalized Laplacian eigenvalues of the distance-based graphs relative to the correlation and learned graphs.

**Figure 5:**
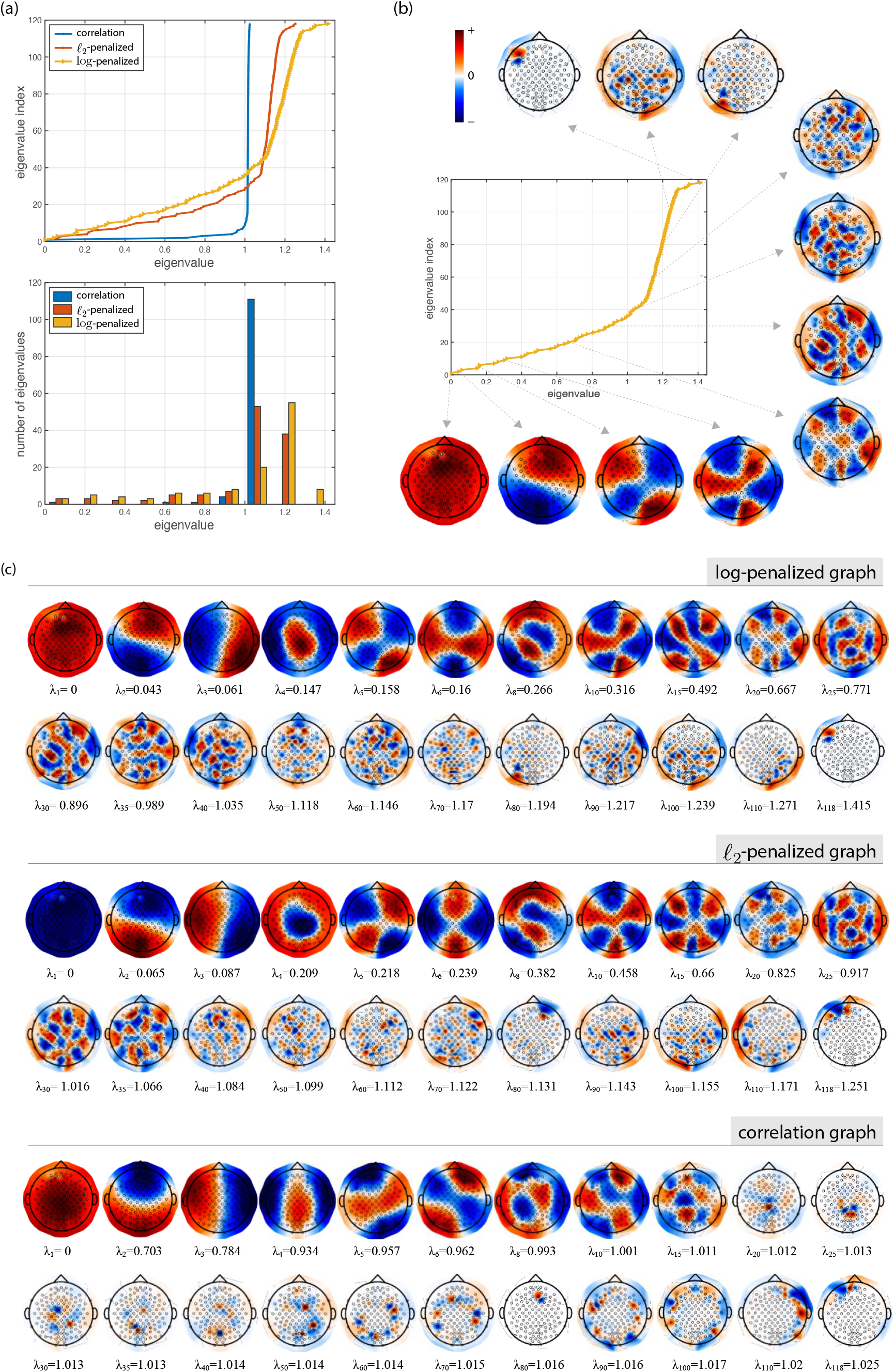
(a) The normalized Laplacian spectra of the log-penalized, *ℓ*_2_-penalized, and correlation graphs of subject aa; the exact eigenvalues are shown on the top whereas the histogram of their distribution is shown at the bottom. (b) A representative set of eigenmaps of the log-penalized graph of subject aa, selected across the graph’s spectral range. (c) A selected subset of eigenmodes of the log-penalized, *ℓ*_2_-penalized, and correlation graphs of subject aa; eigenmaps associated with the same eigenvalue indices across the three graphs are displayed.

Figure 5(b) illustrates eigenmaps associated to a representative set of normalized Laplacian eigenvalues of the log-penalized graph. The first eigenmap is almost evenly distributed over all the graph vertices and given that TV(**u**_1_) = *λ*_1_ = 0 there is no notable spatial variation. In the next eigenmaps, the increase in spatial variability is proportional to the increase in graph frequencies. The last eigenmap is highly localized, which is in line with normalized Laplacian matrices characteristics that manifest localized patterns of spatial variability in high frequencies. Figure 5(c) shows several of the eigenmaps and their corresponding eigenvalues for the three studied graphs for subject aa. The eigenmaps of the learned graphs capture a wider range of variability compared to the correlation graph, many eigenmaps of which manifest spatial patterns with a similar variability corresponding to a spectral value around one. The complete set of eigenmaps of the three studied graphs and the two distance-based graphs for subject aa are shown in supplementary material Figures S4-S8.

Aside from studying the structure and properties of the learned graphs, it is also intuitive to quantify their structure in relation to graph signals. Given that smoothness is a main term in the objective function of the leveraged learning methods, as a first step, we validated the degree to which graph signals of the test-set data are represented as smooth signals on the graphs that were learned using the training-set data. For each subject, for each learned graph, and for each EEG trial, we computed the mean TV of the set of graph signals within that trial (average over 200 graph signals); in particular, using each graph’s **L** matrix, we computed the trace term in (14), and divided the resulting value by the number of graph signals, 200, to obtain an average TV value for each trial. We performed this analysis not only for the learned graphs, but also for correlation graph for comparison. Moreover, to verify the efficiency of the learning process in resulting in a graph on which graph signals of the test-set data are smooth compared to that on a null graph, we generated surrogate Laplacian matrices from the learned graphs (also the correlation graph) by randomly shuffling the edges, while maintaining matrix symmetry; a new surrogate Laplacian matrix was generated for each trial. With this approach, firstly, we generate random graphs with the same sparsity as that of the learned graphs (but complete graphs in the case of the correlation graph), and secondly, we retain the edge weight distribution. The results are presented in Figure 6(a). EEG graph signals show a substantially lower TV on the learned graphs compared to, on the one hand, the correlation graphs, and on the other hand, their surrogate graphs, across all subjects. This reflects the efficacy of the learning process in providing a domain on which EEG signals are smooth. As a second step, we validated whether the learned graph for each subject provides a better substrate on which the subject EEG graph signals are most spatially smooth. The results are presented in Figure 6(b). In all five subjects, EEG graph signals have lowest TV on the graph learned using the subject data compared to graphs learned from other subjects data; see entries on the main diagonal for the learned graphs. However, this pattern is not seen on the correlation graphs, for which the graph of subject al provides the substrate on which 3 out 5 subjects’ data are seen as most smooth—see second colum in the third matrix— and that of subject aw provides a substrate on which the highest values of TV (least smoothness) are seen not only for the signals of subject aw but also for the signals of the other four subjects—see fourth column in the third matrix. The TV of the training-set data on the learned graphs is presented in Figure S9 in the supplementary material.

**Figure 6:**
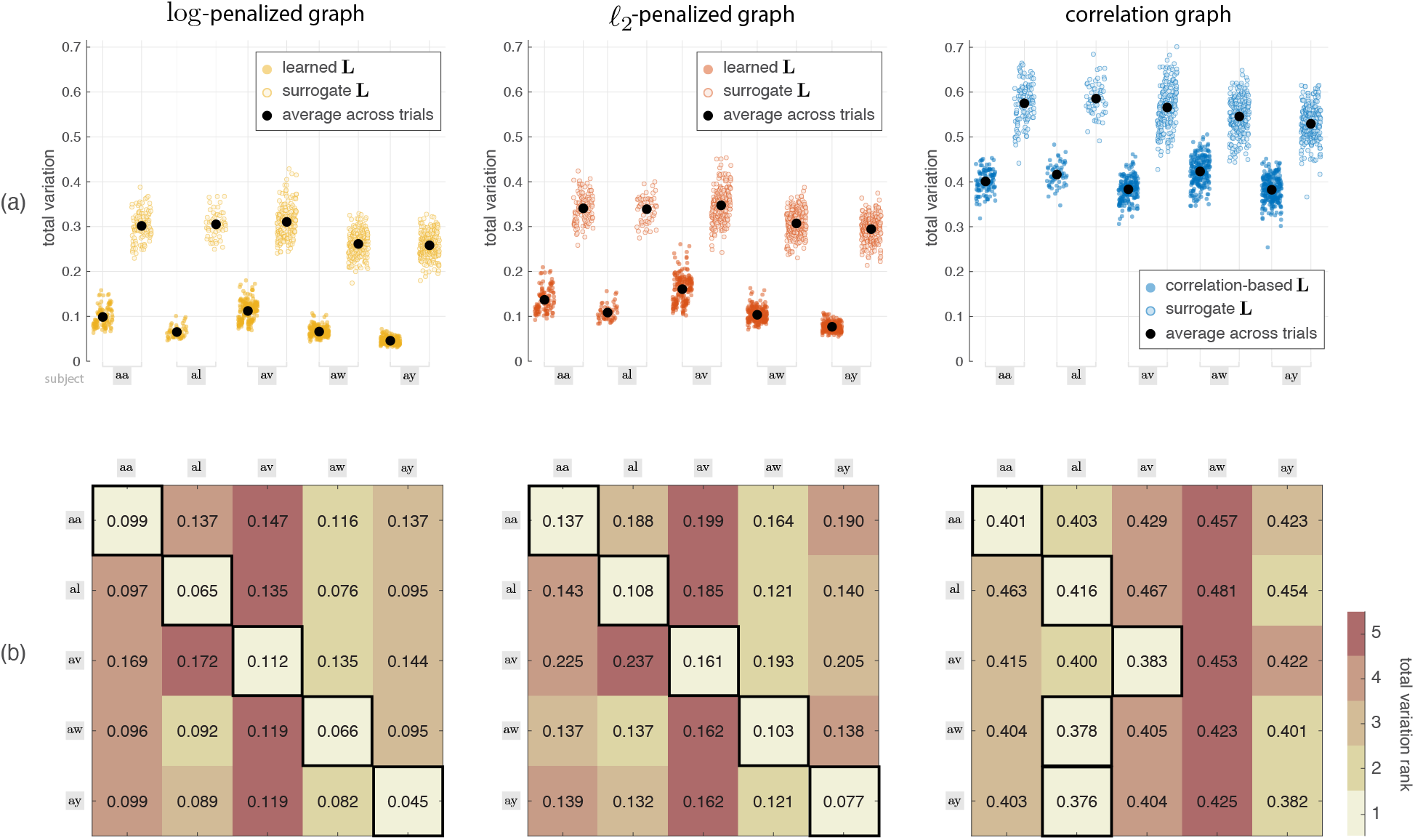
Validation of the extent of smoothness of the EEG graph signals on the learned graphs and correlation graphs; TV values are shown, the inverse of which is a measure of smoothness. (a) TV of graph signals on the learned/correlation graphs as well as surrogate graphs in all subjects. (b) The mean TV of each subject’s graph signals on its learned/correlation graph and on the other subjects’ graphs; Each row in a matrix—ranked via color-code—represents the mean TV of signals of a single subject on the graph of the different subjects; e.g., the element on the third row and fourth column is the mean TV of signals of subject av on the learned graph of subject aw, whereas the third diagonal element is the mean TV of signals of subject av on the learned graph of the same subject.

The WZC of the normalized Laplacian eigenvectors of the three studied graphs is shown in Figure 7(a). Spatial variability of eigenvectors generally increases by increasing the eigenvalue indices. WZC gradually increases along the spectrum in the learned graphs, especially in the log-penalized one, whereas in the correlation graph, it sharply increases in the initial eigenvalue indices, and then only minimally changes in the remainder of the spectrum. Figure 7(b) shows the relation between the WZC of the eigenvectors and their related eigenvalues. The eigenvalues of the learned graphs provide a more uniform sampling of the spectral range whereas those of the correlation graph are cluttered within the upper end of the spectrum. Moreover, the eigenvalues of the learned graphs show a clear relation to the WZC of their corresponding eigenvectors, a consistent second order polynomial trend, which is not observed in those of the correlation graph. Figure S10 in the supplementary material provides a comparison between the WZC of the eigenvectors of distance-based graphs relative to that of the correlation and learned graphs.

**Figure 7:**
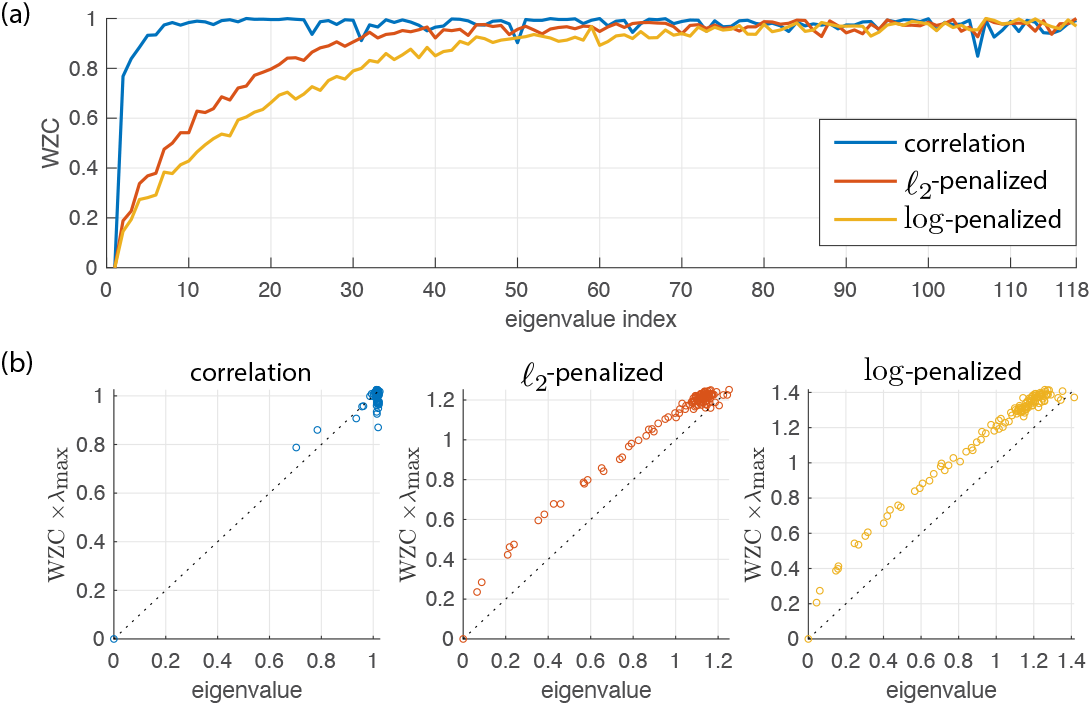
(a) Weighted zero crossings measure for the eigenvectors of the studied graphs for subject aa. (b) Relation between the WZC measure of the eigenvectors and their corresponding eigenvalues.

Figure 8 shows the cumulative energy spectra of test-set data graph signals on the correlation and learned graphs build using the training-set data of subject aa. Results are also presented on the surrogate graphs, constructed using the same approach described in TV analysis for Figure 6. In both learned graphs, a substantial proportion of the energy of EEG graph signals is concentrated in the lower-end of the spectrum, with only a mere fraction of lowest frequency eigenvectors (approximately 17%) capturing most of the signal energy (approximately 90%). The energy spectra sharply increase in the initial eigenvalues of the log-penalized and *ℓ*_2_-penalized graphs, the increasing rate of which decreases in the remainder of their spectra. For the correlation graph, as in the two learned graphs, approximately 70% of signal energy is captured by only the second to seventh eigenvectors, however, the covered spectral range substantially differs between the two, where for the correlation graph almost 95% of the graph’s spectrum is covered by this amount of energy, whereas in contrast, only 25% and 35% of the spectrum is covered for the log-penalized and *ℓ*_2_-penalized graphs, respectively. For the surrogate graphs, the energy almost uniformly increases as a function of the number of spectral indices; i.e., an equal amount of energy is captured by each eigenvector, a characteristic that is strongly reminiscent of the spectral structure of white noise. In Figure S11 in the supplementary material, cumulative energy spectra profiles of both the test-set and training-set data are presented, using not only functional and surrogate graphs as in Figure 8, but also for distance graphs, and across the five subjects.

**Figure 8:**
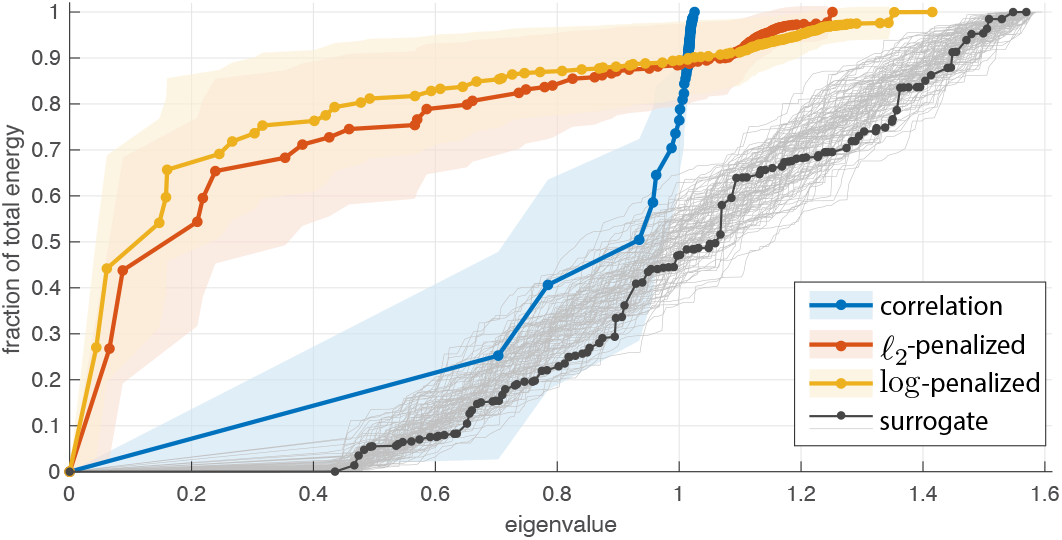
Cumulative energy spectra of EEG graph signals from subject aa on different graphs. Aside from the three studied graphs, results are presented on 100 surrogate graphs; given that each surrogate graph has a unique spectrum, results are presented as individual curves rather being averaged; the eigenvalues of a representative surrogate graph are marked.

In order to investigate the performance of the proposed method in MI-classification, five different sets of the GFT coefficients were utilized in the first experiment. The first set consisted of the entire set of GFT coefficients, denoted all frequencies (AF). Similar to prior works on the application of GFT on brain imaging data (Medaglia et al., 2018; Jafadideh and Asl, 2022), three additional sets of GFT coefficients were defined by dividing the spectrum into three equal frequency bands, denoted low (LF), medium (MF) and high (HF) frequencies. Inspired by (Cattai et al., 2021), a fifth subset was defined via the union of the LF and HF subsets, denoted LF+HF. These five sets of GFT coefficients were then used as inputs to the FKT to derive a discriminative matrix *W* for each set. Consequently, features for classification were extracted by computing the logarithm of variance of the projected GFT coefficients on *W*.

Table 2 presents classification accuracies using the three different graphs for each individual subject and also on average across subjects. Using the LF GFT coefficients resulted in substantially higher classification accuracy compared to using the MF or HF components, in all subjects as well as on average across subjects. In comparison to AF and LF+HF, the LF coefficients provided the highest accuracy in subjects aa, and ay in all three graphs. In subject aw, LF showed highest accuracy for log-penalized and correlation graphs, whereas it was outperformed by AF for *ℓ*_2_-penalized graph. In subject al, LF showed similar results as AF and LF+HF for log-penalized and correlation graphs, whereas it was outperformed by AF and LF+HF for *ℓ*_2_-penalized graph. In subject av, best results were obtained using the LF+HF components, in all three graphs. Overall, the lowest classification accuracies in both learned graphs are for subject av, corroborating the highest mean TV (least smoothness) of this subject’s graph signals on its learned graphs as shown in Figure 6.

**Table 2:** Classification accuracies (in %) for the three different brain graph designs on the test-set data for each subject, and on average across subjects, in five different graph frequency band settings.

| log-penalized | aa | al | av | aw | ay | Mean±std |
| --- | --- | --- | --- | --- | --- | --- |
| AF | 74.11 | 100 | 70.41 | 90.18 | 74.21 | 81.78 ± 12.73 |
| LF | 86.61 | 100 | 70.92 | 91.96 | 84.92 | **86.88 ± 10.68** |
| MF | 55.36 | 66.07 | 52.55 | 63.39 | 54.36 | 58.35 ± 5.99 |
| HF | 51.78 | 80.36 | 43.88 | 59.37 | 47.22 | 56.62 ± 14.53 |
| LF+HF | 74.11 | 100 | 71.43 | 90.62 | 75.79 | 82.39 ± 12.35 |

| $\ell_2$-penalized | aa | al | av | aw | ay | Mean±std |
| --- | --- | --- | --- | --- | --- | --- |
| AF | 69.64 | 100 | 70.41 | 89.73 | 72.22 | 80.40 ± 13.73 |
| LF | 87.5 | 98.21 | 70.41 | 83.03 | 88.49 | **85.53 ± 10.10** |
| MF | 54.46 | 67.86 | 53.06 | 62.05 | 55.16 | 58.52 ± 6.27 |
| HF | 54.46 | 80.36 | 44.39 | 55.80 | 51.19 | 57.24 ± 13.66 |
| LF+HF | 78.57 | 100 | 72.45 | 86.61 | 78.57 | 83.24 ± 10.63 |

| correlation | aa | al | av | aw | ay | Mean±std |
| --- | --- | --- | --- | --- | --- | --- |
| AF | 70.53 | 100 | 70.92 | 88.39 | 72.22 | 80.41 ± 13.25 |
| LF | 86.61 | 100 | 68.88 | 94.2 | 87.7 | **87.47 ± 11.71** |
| MF | 52.68 | 78.57 | 54.59 | 54.91 | 53.97 | 58.94 ± 11 |
| HF | 55.36 | 71.43 | 56.12 | 51.34 | 48.41 | 56.53 ± 8.89 |
| LF+HF | 66.96 | 100 | 71.43 | 91.96 | 84.13 | 82.9 ± 13.8 |

We used feature selection to determine an optimal subset of the GFT coefficients that provide the most discriminative features for classification; cf. Section 2.5.2. The logarithm of variance of the GFT coefficients was used as input to feature selection. Figure 9 illustrates the scores of graph frequencies in the log-penalized graph for each subject and on average across subjects. The lowest one-third eigenvalue indices attained substantially higher scores than the rest of the spectrum, corroborating results presented in Figure 8 and in Table 2 that show EEG signal energy is largely captured by eigenvectors within the LF range. Therefore, we only used selected features from this sub-band as the most effective harmonics for each subject, classification accuracy for which are presented in Table 3. To evaluate the effectiveness of using the FKT, results of classification using GFT coefficients (without FKT) are also provided in Table 3. The direct use of the GFT coefficients is prone to over-fitting due to the small size of the training samples in comparison to the dimension of the feature vectors, especially in the subjects with small training sets. Therefore, a subset of GFT coefficients as determined by the feature selection step were fed into the classifier. Comparing these results with those in Table 2 reveals that for all three graphs, the performance in two of the subjects—aa and ay for log-penalized and correlation, av and aw for *ℓ*_2_-penalized—enhanced when selected features from LF were used compared to when all LF coefficients were used, which has in turn resulted in better overall accuracy. Moreover, the results suggest that using FKT notably improves classification accuracy compared to directly using the GFT coefficients. That is, mapping the GFT coefficients onto the subspace provided by FKT results in features that better discriminate the two MI classes, thanks to accounting for the difference in the covariance structure (Cohen, 2022) of the GFT coefficients of the two classes. Figure 10(a) shows the obtained FKT filters for the log-penalized graph of subject aa. The most discriminative coefficients can be inferred from the absolute value difference between two filters, shown in Figure 10(b); the corresponding eigenmaps of the six top-ranked indices are shown in Figure 10(c), whereas, for comparison, those for the *ℓ*_2_-penalized and correlation graphs are shown in Figures 10(d) and (e), respectively. The FKT filters for the other four subjects are presented in Figure S12 in the supplementary material. Overall, the best average accuracy in Table 3 was obtained in the proposed method by using the log-penalized graph learning approach. Related results on using distance-based graphs are presented in Tables S1S2 in the supplementary material.

**Table 3:** Classification accuracies (in %) when directly using the GFT coeffi-cients vs the proposed method, wherein GFT coefficients are subjected to FKT. In both settings, features are selected from the LF sub-band of the graph spectra.

| GFT | aa | al | av | aw | ay | Mean±std |
| --- | --- | --- | --- | --- | --- | --- |
| log-penalized | 61.61 | 87.5 | 60.71 | 70.53 | 81.75 | 72.42 ± 11.96 |
| $\ell_2$-penalized | 65.18 | 91.07 | 57.65 | 67.86 | 66.27 | 69.61 ± 12.62 |
| correlation | 68.75 | 96.43 | 62.75 | 73.21 | 68.65 | 73.96 ± 13.1 |
| Proposed | aa | al | av | aw | ay | Mean±std |
| log-penalized | 87.5 | 100 | 70.92 | 91.96 | 92.86 | **88.65 ± 10.88** |
| $\ell_2$-penalized | 87.5 | 98.21 | 72.96 | 84.82 | 88.49 | 86.4 ± 9.06 |
| correlation | 90.18 | 100 | 68.88 | 94.2 | 88.89 | 88.43 ± 11.75 |

**Figure 9:**
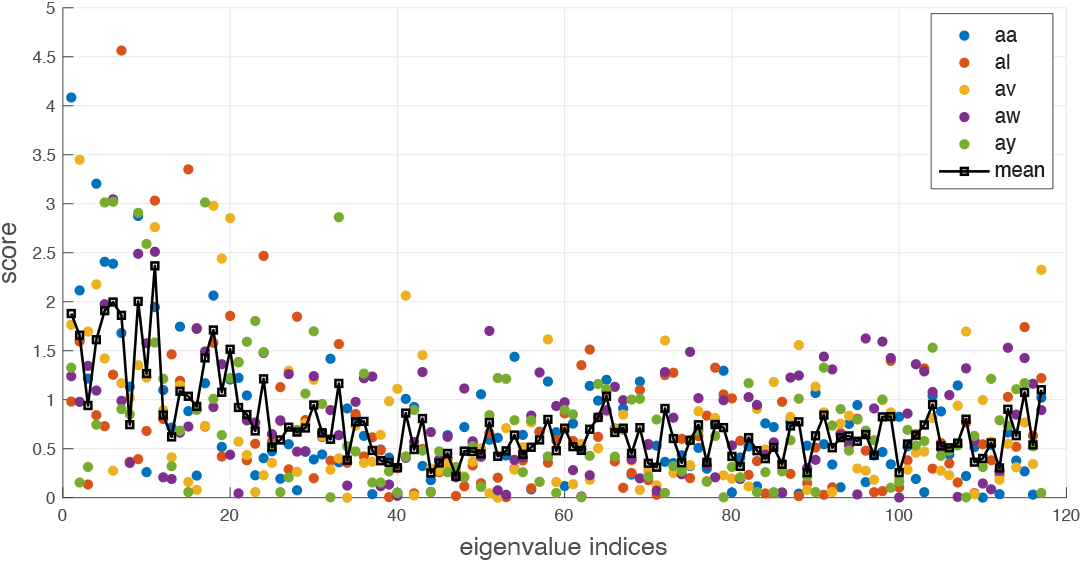
Scores of the graph frequencies in the log-penalized graph for each individual subject and on average across subjects.

**Figure 10:**
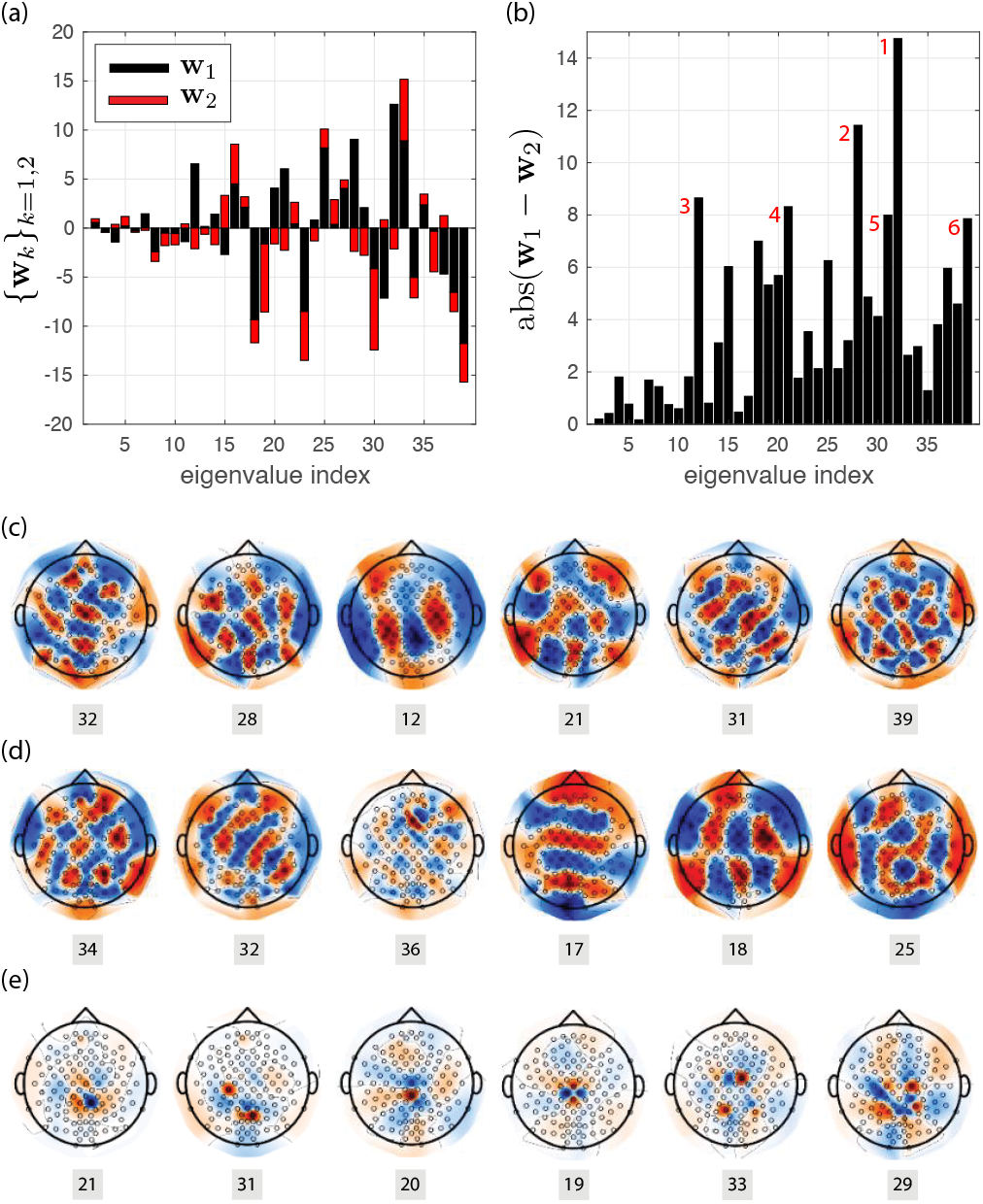
(a) FKT filters for the log-penalized graph of subject aa using the lower end of the spectrum. (b) Absolute value difference between the two filters, indicating the harmonics which provide maximal contribution for discrimination the two classes. (c) The top six eigenmaps based on (b); indices below each map indicate eigenvalue index. (d)-(e) Same as in (c) but for the subject’s *ℓ*_2_-penalized and correlation graphs, respectively; the FKT filters for these two graphs of subject aa are shown in supplementary Figure S12.

Finally, the performance of the proposed method was compared to four recent alternative state-of-the-art methods: a GSP-based method (Georgiadis et al., 2021) that utilizes graph Slepians (Van De Ville et al., 2017), and three methods that utilize different extensions of the FKT (Cherloo et al., 2021; Wang et al., 2020; Hou et al., 2022). The results are presented in Table 4. The proposed method using log-penalized graph learning outperforms the alternative methods, on average across subjects. The method by (Georgiadis et al., 2021) shows the best classification accuracy in subject av, whereas that of (Cherloo et al., 2021) shows the best accuracy in subject aw. In the other three subjects, the proposed method yields higher classification accuracy.

**Table 4:**
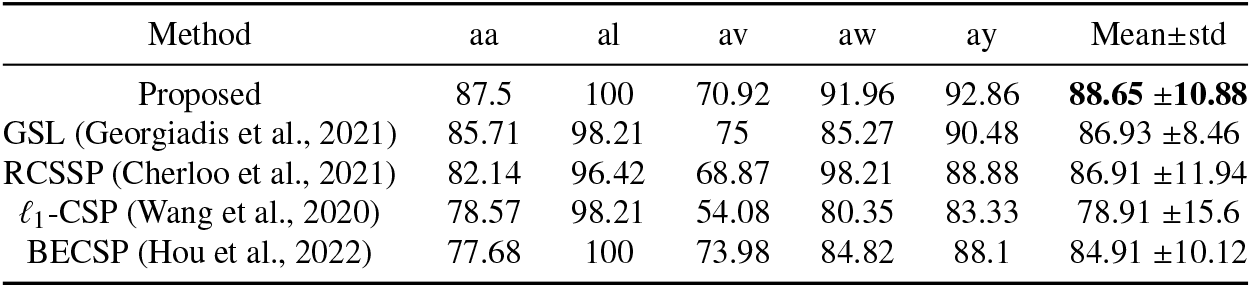
Comparison of classification accuracies (in %) for the proposed method (log-penalized graph, GFT+FKT, selected features from the LF sub-band) and four alternative state-of-the-art methods.

| Method | aa | al | av | aw | ay | Mean±std |
| --- | --- | --- | --- | --- | --- | --- |
| Proposed | 87.5 | 100 | 70.92 | 91.96 | 92.86 | **88.65 ± 10.88** |
| GSL (Georgiadis et al., 2021) | 85.71 | 98.21 | 75 | 85.27 | 90.48 | 86.93 ± 8.46 |
| RCSSP (Cherloo et al., 2021) | 82.14 | 96.42 | 68.87 | 98.21 | 88.88 | 86.91 ± 11.94 |
| $\ell_1$-CSP (Wang et al., 2020) | 78.57 | 98.21 | 54.08 | 80.35 | 83.33 | 78.91 ± 15.6 |
| BECSP (Hou et al., 2022) | 77.68 | 100 | 73.98 | 84.82 | 88.1 | 84.91 ± 10.12 |

## 4. Discussion

Volume conduction results in spatial smearing in EEG electrode measurements (Schaworonkow and Nikulin, 2022), making it of importance to account for underlying spatial structures. A graph model can provide a holistic view of such spatial dependencies. Here we studied three graph models, one based on the Pearson correlation between pairs of electrodes, and two based on capturing the inherent spatial smoothness manifested jointly by the entire set of electrodes via graph learning. For each subject, the studied graphs present a notable difference in their nodal degrees, and their structure in general, showing the added benefit of using subject-specific graphs, which is inline with the body of literature that suggest intersubject variability in EEG spatial patterns manifested for identical tasks (Blankertz et al., 2007; Zhang et al., 2016; Zabicki et al., 2019; Corsi et al., 2020). Learned graphs of subjects aa, al and aw have larger nodal degrees, and edge weights are seen in motor-related areas in the frontal and parietal cortices, areas that are commonly known to become active under MI tasks (Pilgramm et al., 2016; Ogawa et al., 2022). In addition to motor-related areas in the frontal and parietal cortices, the correlation graphs of all subjects also manifest large nodal degrees in the occipital lobe; these patterns can be interpreted as high spatial frequencies (correlated asynchronous activity between electrode pairs), a pattern which is in contrast penalized by the smoothness criterion in the objective function of the graph learning algorithms. The log-penalized graphs are sparser graphs with larger edge weights compared to the *ℓ*_2_-penalized graphs. This sparsity is desirable as it, on the one hand, is a characteristic of realistic networks such as that of the human brain, and on the other hand, reduces the computational burden of algorithms, making the method suitable for online BCI applications.

Given that graph Laplacian eigenvectors form an orthonormal basis for graph signals (Chung, 1997), their broader spatial variability with respect to the graph structure—i.e., range and distribution of eigenvalues, cf. Figure 5—can provide a more effective decomposition when computing a spectral representation of EEG graph signals via the GFT. The WZC quantification of the eigenvectors of the correlation and learned graphs, cf. Figure 7, corroborate visual interpretations made on spatial variability of eigenmaps as shown in Figure 5(c), reflecting the superior capability of the learned graphs over the correlation graph in providing a basis that represents a broader range of spatially varying patterns. The effect of this difference in the eigenbasis can also be seen by inspecting the spectral representation of the associated EEG signals on the graphs. In the correlation graph, a small subset of the first eigenvectors captures a substantial portion of the total signal energy, whereas in the learned graphs, signal energy is distributed across a wider range of eigenvectors. In particular, eigenvalues with high multiplicity around one in the correlation graph spectrum (a high peak at *λ ≃* 1) suggest vertex duplication, in which a new vertex to the graph has an identical connectivity pattern to the duplicated vertex, resulting in vertices with the same connectivity profile (de Lange et al., 2014). This suggests that the connectivity pattern is rather similar across vertices in the correlation graph.

The proposed graph learning and GSP setting enables an intuitive accounting of spatial dependencies observed in EEG data. However, it does not account for temporal dynamics, aside from using the ensemble set of EEG graph signals when performing graph learning. To jointly account for the spatiotemporal features, we leveraged the FKT, which functions on the spectral representation of EEG data as provided by GFT. The proposed FKT-based approach of extracting features from a temporal set of GFT coefficients is in contrast to prior related works (Huang et al., 2018; Preti and Van De Ville, 2019; Bolton and Van De Ville, 2020) wherein the temporal mean or variance of the GFT coefficients is considered as feature, which notably discards the temporal dynamics. The temporal evolution of GFT coefficients of two representative EEG trials and the absolute value of the difference between them are shown in Figure 11. Largest GFT coefficients are located mainly in the lower end of the spectrum, reflecting the relevance of spatial patterns manifested by the eigenvectors within the LF range (cf. Figure 5) in representing a substantial extent of the variance observed in EEG signals. Subtle differences are seen between the two classes. The GFT coefficients are not consistent across time, showing notable variability. By using the FKT, the manifested temporal dynamics are taken into account, mapping the GFT coefficients to a subspace in which the two MI tasks can be effectively differentiated, corroborating related findings on fMRI data (Itani and Thanou, 2021).

**Figure 11:**
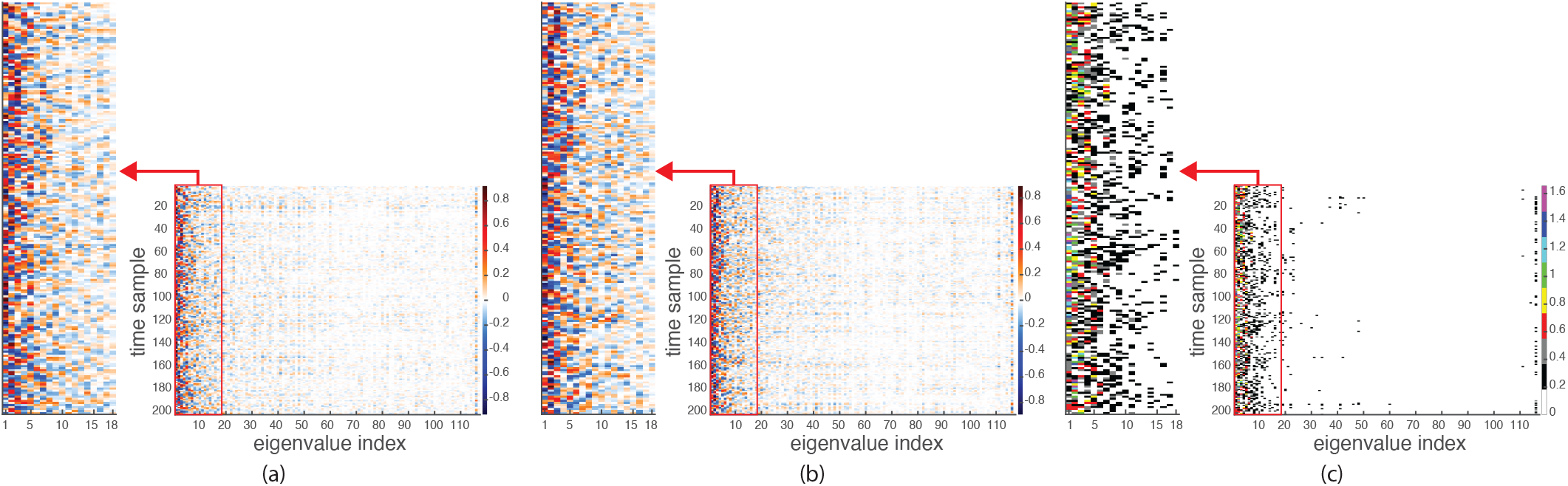
GFT coefficients of the log-penalized graph for a representative trial of the (a) right hand and (b) right foot MI classes of subject aa; each trial consists of 200 time instances, thus, 200 graph signals. (c) The absolute value of the difference between the GFT coefficients of the trials shown in (a) and (b).

### 4.1. Limitations and Outlook

In relation to the definition of brain graph vertices, as in related prior works—e.g. (Saboksayr et al., 2021; Georgiadis et al., 2021; Cattai et al., 2021), we treated each EEG electrode as a graph vertex, thus, directly used EEG electrode recordings after minimal preprocessing, without utilizing source reconstruction (Farahibozorg et al., 2018). For the application considered in study, i.e. MI task decoding, source reconstruction is not required, and in fact, it is favorable to not use as it increases computational speed within a BCI setting, and also prevents undesired spatial smoothing that results from source-reconstruction. Nevertheless, in other applications, in particular, those of cognitive neuroscience, it is beneficial to define a brain graph wherein brain regions derived from a template atlas are treated as vertices, e.g. as in (Rué-Queralt et al., 2021; Glomb et al., 2020a), which would reduce spurious connectivity between vertices (Van de Steen et al., 2019); this would require using a dataset that includes suitable structural or diffusion MRI data to perform source reconstruction, which may however not be available in some datasets. The graph learning method proposed in this paper can be extended to use source-reconstructed EEG data; regularization parameters—*α* and *β*—used in the optimization terms may need tuning to account for the difference in the nature of nodes, and as such, may for example be set by linking them to desired graph sparsity (Kalofolias and Perraudin, 2017). By using source reconstruction, the resulting graphs and eigenmaps can be, firstly, more easily interpreted in relation to underlying structure, and secondly, matched and subsequently averaged or compared across subjects, and used for applications such as fingerprinting (Gao et al., 2021; Sareen et al., 2021; Griffa et al., 2022).

The spatial dependency between brain activity at neighboring electrodes—resulting from the smearing effect that is inherent in EEG—suggests the need to take into account the spatial configuration of electrodes, a topic of continued research interest (Iivanainen et al., 2021; Conrad et al., 2020). The structure of the functional graphs studied in this paper—both for the correlation and the learned graphs—is affected by the geometric configuration of electrodes. Here we studied differences in harmonic basis derived from functional graphs to that of distance-based graphs via quantifying differences in signal energy profiles or MI-task decoding performance. To study the effect of distance, we assessed the degree to which geometric configuration of electrodes is sufficient to spatially decompose EEG data using distance-based graphs, the results of which are presented in the supplementary material. In particular, MI-decoding results showed a relatively inferior performance for distance-based graphs relative to learned graphs (cf. Tables S1-S2), indicating the effectiveness of inferring functional and structural relation from EEG maps (Pang et al., 2022; Suärez et al., 2020). Future work is necessary to provide an indepth comparison between EEG harmonics of different nature, with the broader goal of obtaining a harmonic basis that can disentangle signal contribution associated to volume conduction from that associated to pure functional relation between channels.

A number of alternative feature extraction strategies may be considered in future work. Firstly, without resorting to GSP, the resulting learned graphs may be readily used for fingerprinting (Gao et al., 2021) or to extract features that characterize brain functional structure, via principles from either graph theory, e.g. as in (Sporns, 2022; Yun and Kim, 2021), or spectral graph theory, e.g. as in (Fan and Chou, 2018; Wachinger et al., 2016; Maghsadhagh et al., 2021). Secondly, for MI task decoding in particular, given that the frontal and parietal cortices are more involved, by choosing a subset of electrodes—e.g. as in (Georgiadis et al., 2021; Cherloo et al., 2021), graph learning can be localized to these regions of interest. Furthermore, it may be beneficial to extract coarse features at the resolution of spectral graph frequency bands rather than GFT coefficients, e.g. as in (Behjat et al., 2015, 2021), as this can be necessary in deriving compact, interpretable EEG-based clinical biomarkers (Mussigmann et al., 2022). The challenge to this aim would be to use an appropriate choice of graph filter banks Shuman (2020); Isufi et al. (2022) based on the graph spectra (Shuman et al., 2015) and the energy spectral density (Behjat et al., 2016) of EEG maps on learned graphs.

An intriguing avenue for future research is to study the relationship between harmonic basis of EEG learned graphs and EEG microstates (Michel and Koenig, 2018; Van De Ville et al., 2017; Mishra et al., 2020)—short periods during which the topographies of EEG maps remain quasi-stable. In particular, given the objective of the learning process to derive a manifold on which observed EEG maps are smooth, subject-specific EEG microstates, which exhibit smooth topographies, may naturally arise as a subset of, or linear combination of, the lowest frequency components of the learned basis. Another avenue of research is to study the potential of EEG learned graphs as a backbone on which spatial filtering of EEG maps can be performed—e.g. for interpolation of missing channels (Banville et al., 2022; Svantesson et al., 2021)—via spectral graph diffusion filtering schemes (Abramian et al., 2021; Tarun et al., 2020). Finally, the proposed EEG-based graph learning and spectral representation via GSP can be readily extended to other data modalities, in particular, fMRI (Itani and Thanou, 2021; Preti and Van De Ville, 2019), near-infrared spectroscopy (Petrantonakis and Kompatsiaris, 2018), or Magnetoencephalography (Tewarie et al., 2019; Sareen et al., 2021).

## 5. Conclusions

We proposed a GSP-based method on learned graphs to extract spatial signal information from EEG data. The applicability of the method was validated within the setting of classifying EEG motor imagery tasks. We treated acquired EEG data at each time-point as a spatial signal that resides on the vertices of three different, subject-specific functional graphs, in particular, two of which were learned from the data. Our analysis showed that imagined motor activities are generally spatially smooth on the learned graphs, and can thus be effectively represented by using only a subset of their graph frequency components. Furthermore, we showed that temporal dynamics manifested in EEG signals can be captured by using the FK transformation, resulting in a discriminative subspace that can better separate motor imagery classes. Classification results showed the superior performance of the proposed method compared to four related alternative methods, indicating the benefit of extracting features via integrating spatial and temporal characteristics of EEG signals within a GSP setting. We believe that the proposed methodology for spectral graph representation of electrophysiological brain data holds the potential to enhance our understanding of brain functional organization in a wide range of applications in neuroinformatics.

## Appendix 1

**Proof of equality (7)**

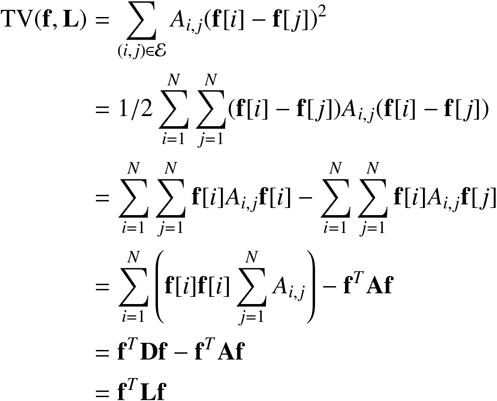

## Appendix 2

**Alternative formulation of TV(f**, L)

An alternative formulation of TV(**f**, *ℒ*) in the form of sum of differences between neighboring vertices—similar to that given in (6) for TV(**f, L**)—can be obtained as:

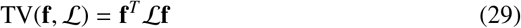

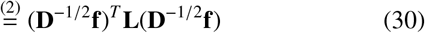

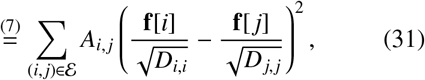

showing the strong similarity between TV(**f, L**) and TV(**f**, *ℒ*), both quantifying the degree of variation in the signal value at elements connected via edge in the graph.

## Appendix 3

**Proof of equality (15)**

Here we reproduce the proof given in (Kalofolias, 2016), adapted to the notations used in the present paper.

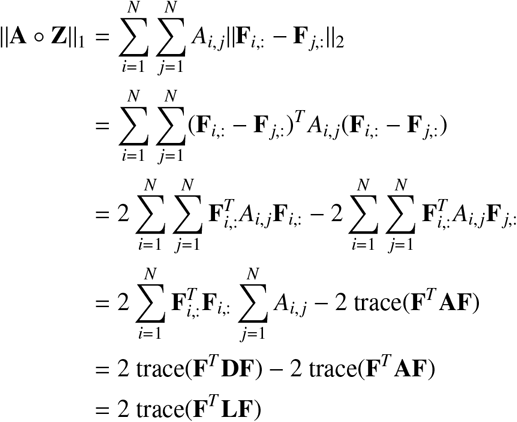

## Appendix 4

**Relation between FKT filters obtained on EEG GFT coe**ffi**cients to those obtained on EEG maps**

In the feature extraction method proposed in this paper, we applied the FKT on the GFT coefficients of EEG maps to derive filters that can discriminate between two classes. In this section, we investigate the mathematical relation between FKT filters obtained via two different approaches: i) using the GFT coefficient of the EEG trials (proposed method), ii) using EEG maps themselves as input to the FKT.

The first approach is thoroughly explained in Section 2.4. Here we derive the relation for the FKT filters using the second approach—commonly referred to within the literature as *Common Spatial Patterns* (Ramoser et al., 2000)—to *those obtained using the first approach; we use the ^∨^ notation to differentiate variables of the same nature in the second approach to those of the first approach. Let* 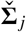 denote the across-trial ensemble averaged covariance matrix of the EEG maps of class *j*, defined as:

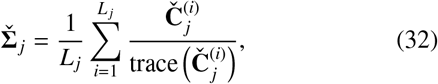

where 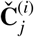 denotes the covariance matrix of the EEG maps of trial *i* of class *j*, obtained as 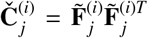. By invoking (4), 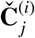 and 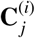 given in (21), can be related as:

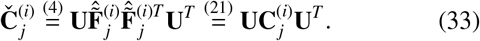

Moreover, invoking the invariance property of the trace under cyclic permutations—i.e., trace(*ABC*) = trace(*CAB*)—and **U**^*T*^**U** = *I*, from (33) we have trace 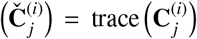. Noting this equality between traces, inserting (33) in (32) gives 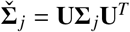, and therefore, 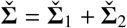 is related to ∑ as:

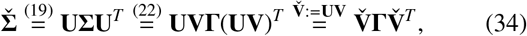

where 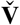 denotes the eigenvectors of 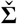. (34) shows that 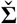 and ∑ share the same eigenvalues and, intuitively, that their eigenvectors are related via the GFT—i.e., 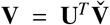, that is, the eigenvectors of ∑ are the GFT of the corresponding eigenvectors of 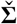. Invoking and using the definition of the whitening matrix given in (23), the relation between whitening matrices is obtained as:

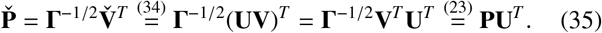

Invoking and using the definition of **S**_1_ as given in (24), **Š**_1_ is obtained as:

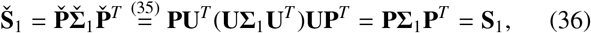

and similarly, we have **Š**_2_ = **S**_2_. Therefore, the eigenvectors of **Š**_1_ and **Š**_2_ are the same as the eigenvectors of **S**_1_ and **S**_2_, i.e. **B** given in (25). Finally, by invoking and using the definition of FKT filters given in (27), the relation between the FKT filters is obtained as:

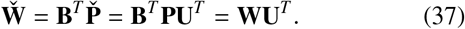

Noting that the FKT filters are given as rows of matrices **W** and 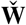, (37) shows that the FKT filters obtained on the GFT coefficients of EEG maps can be equivalently obtained by applying GFT to FKT filters obtained on the EEG maps, a very intuitive relation between the filters from the two approaches.

## Data and code availability

The dataset used in this study is open access and can be downloaded from https://www.bbci.de/competition/iii/. An implementation of the methods proposed in this work will be made available as a MATLAB package on GitHub at https://github.com/aitchbi/.

## Acknowledgments

Data used in this work were provided as part of the BCI Competition III, organized by the Berlin Brain-Computer Interface team https://www.bbci.de/. The authors are grateful to S. Itani and D. Thanou for sharing the code related to the method in (Itani and Thanou, 2021). H. Behjat was supported by the Swedish Research Council under Grant 2018-06689. A preliminary version of this work has been presented in (Miri et al., 2022).

---

**Figure S1:**
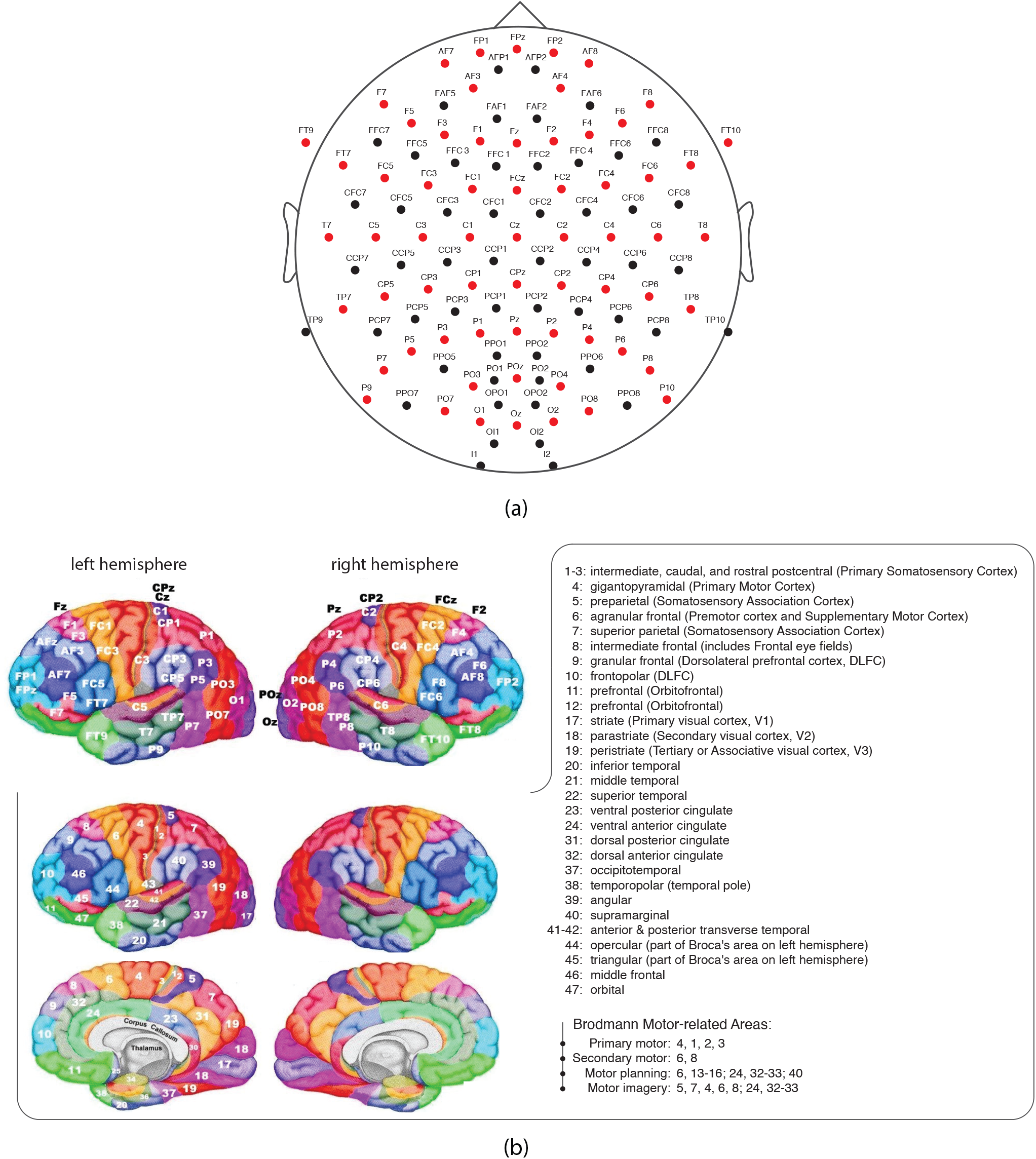
(a) Arrangement of EEG electrodes. The electrodes are labelled, wherein the letters correspond to cortical locations: frontal (F), temporal (T), central (C), parietal (P), occipital (O), Inion (I); combinations of letters indicate intermediate locations. Odd and even numbers within the electrode labels indicate the left and right hemispheres, respectively. The letter ‘z’ (meaning zero) indicates the midline. (b) The position of a subset of the electrodes (64/118, those marked as red in (a)) is shown on the cerebral cortex (top left), in particular, relative to Brodmann area centers (bottom left); the surface maps were adopted from http://www.brainm.com/software/pubs/dg/BA_10-20_ROI_Talairach/nearesteeg.html.

**Figure S2:**
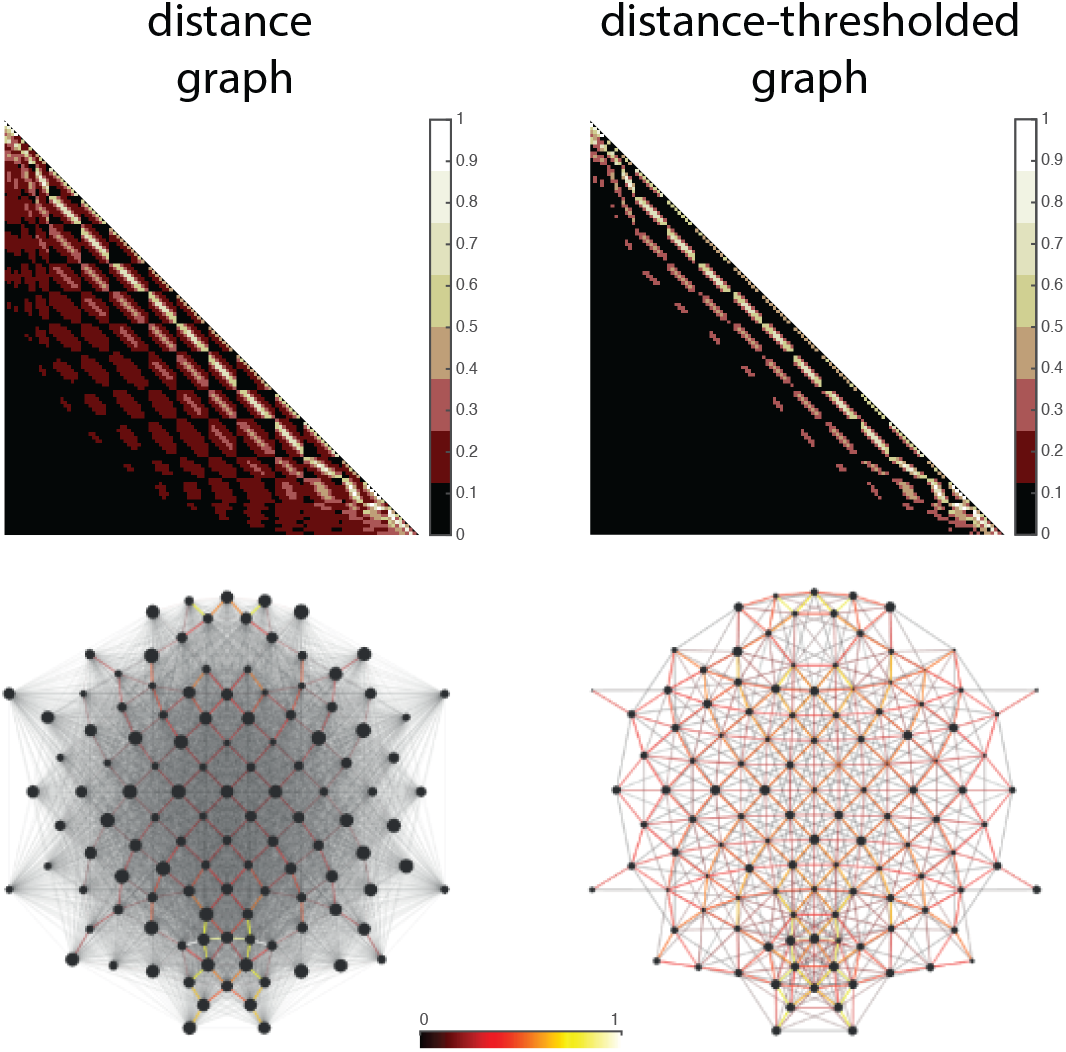
Schematic representation of the distance (left, bottom row) and the distance-thresholded (right, bottom row) graphs with their corresponding adjacency matrices (top row). In the distance-thresholded graph, twelve percent of the graph edges with the largest weights were preserved and the remained edges were removed. This threshold was selected according to the mean of the connection density in the log-penalized graph. Edge widths and colors reflect edge weights, and vertex sizes reflect nodal degrees. For better visualization, only the top 50% of edges that have the largest weights are shown in the distance graph.

**Figure S3:**
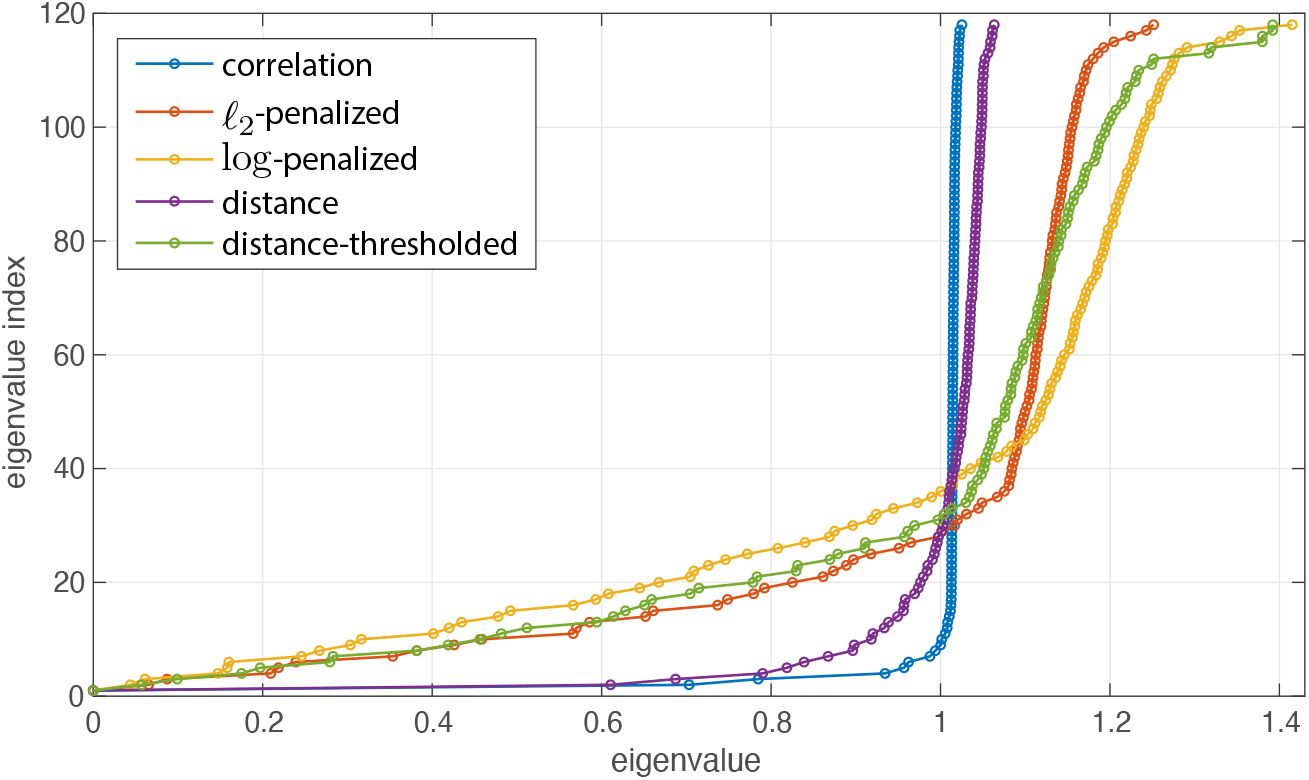
The normalized Laplacian spectra of the three studied graphs as well as the distance-based graphs of subject aa. The distribution of the eigenvalues of the distance graph, which is a complete graph, is similar to the correlation graph. On the other hand, the eigenvalues of the distance-thresholded graph have a similar trend to the eigenvalues of the learned graphs.

**Figure S4:**
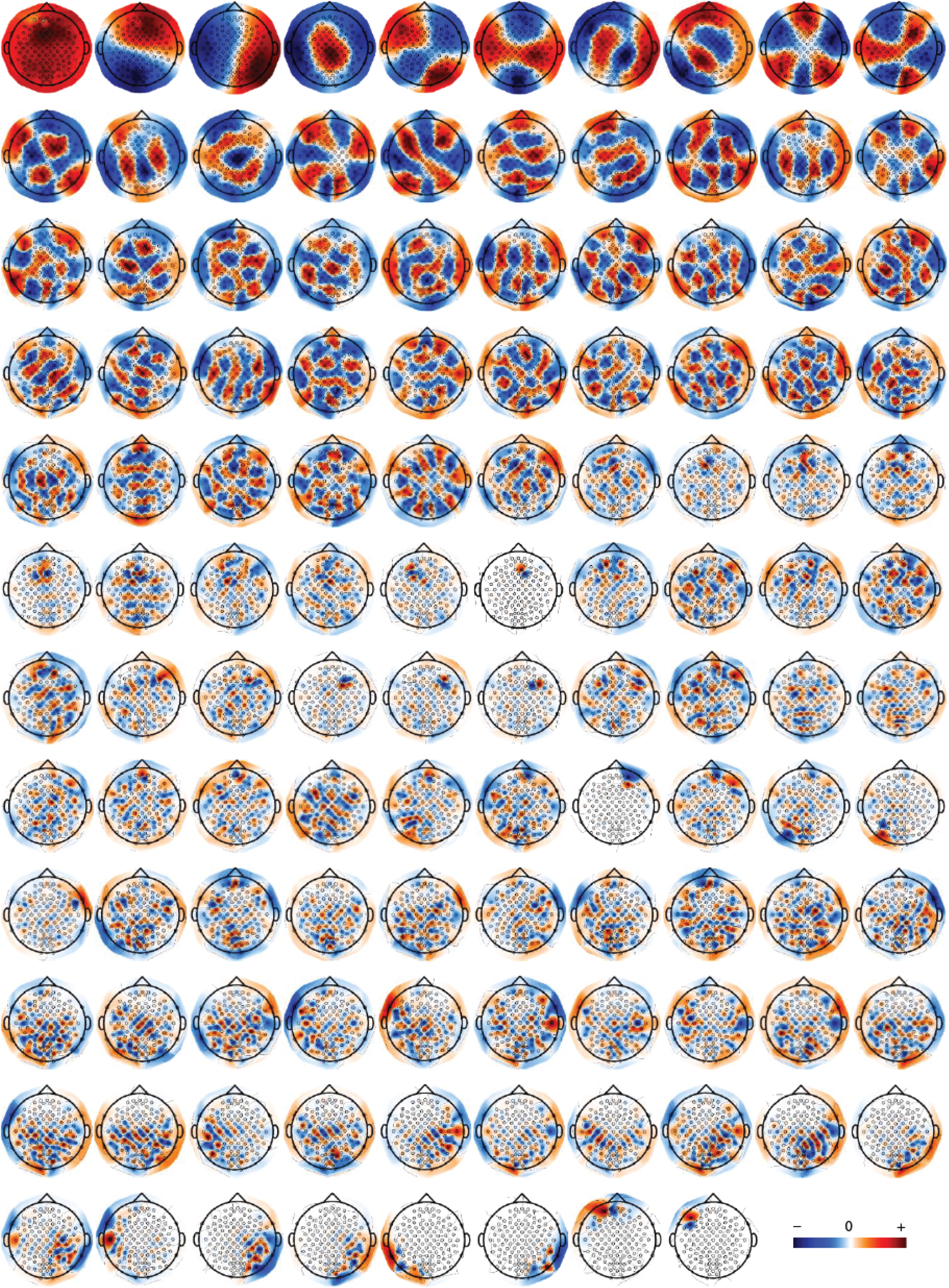
The Laplacian eigenmaps of the log-penalized graph for subject aa.

**Figure S5:**
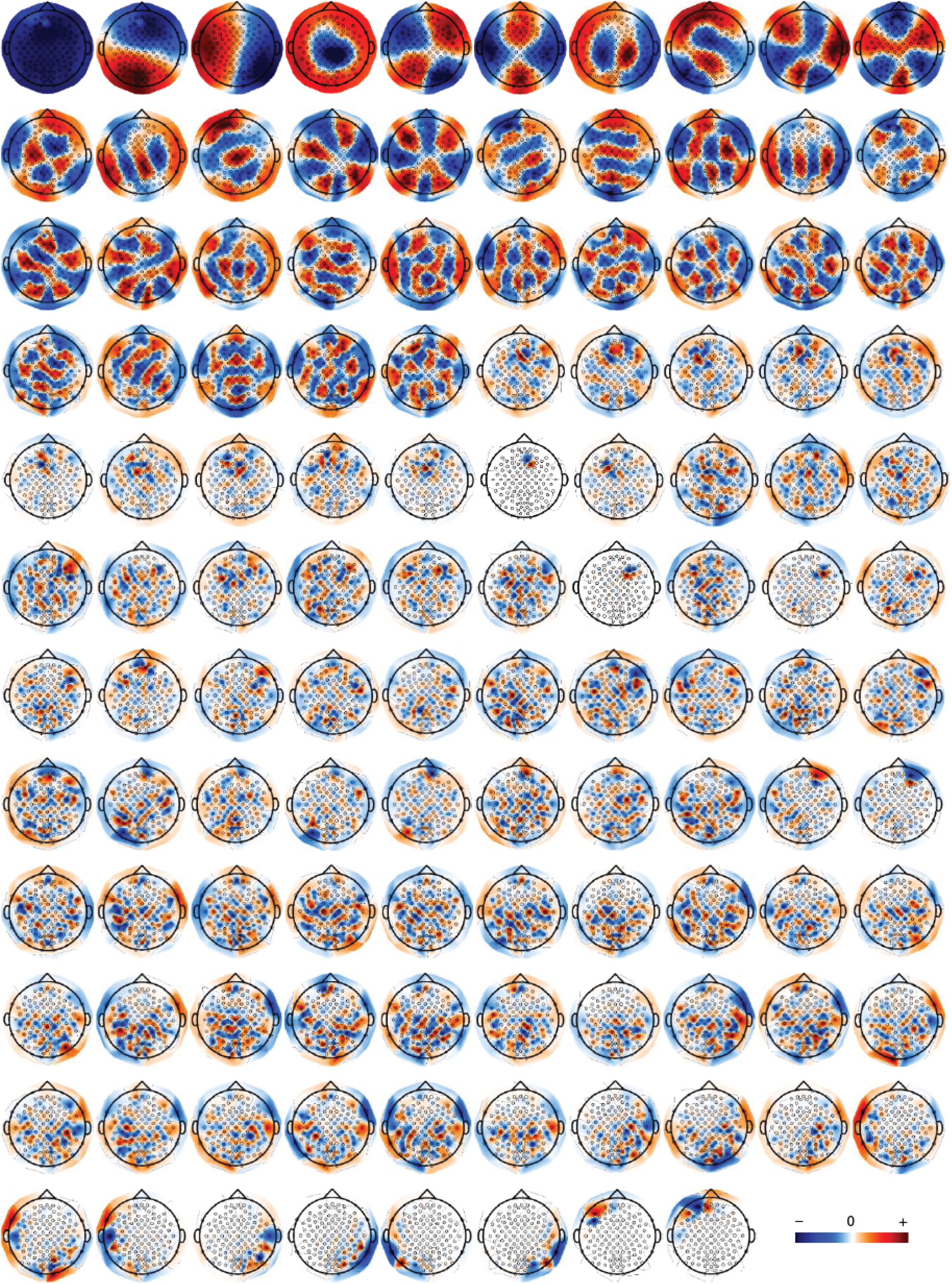
The Laplacian eigenmaps of the *ℓ*_2_-penalized graph for subject aa.

**Figure S6:**
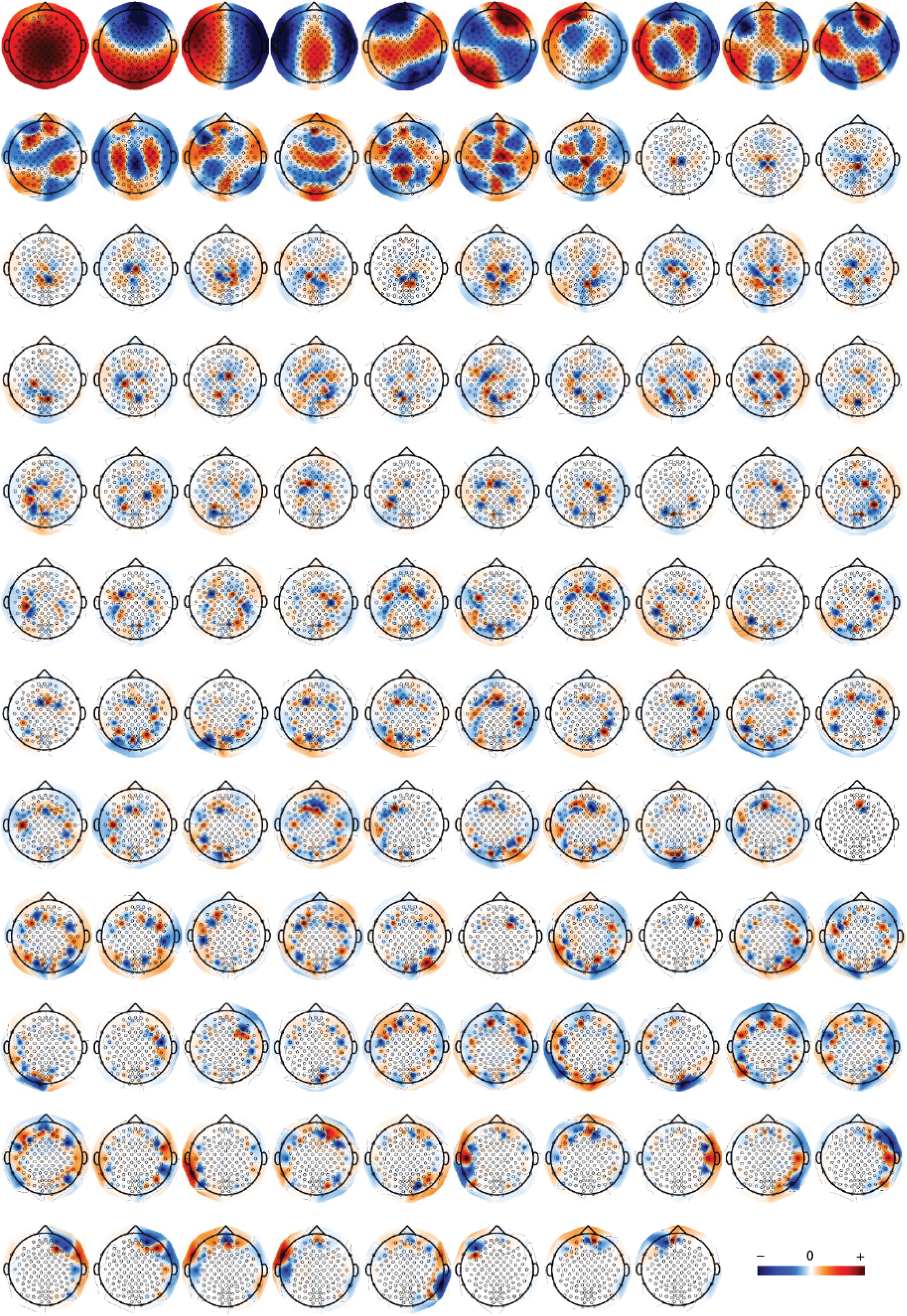
The Laplacian eigenmaps of the correlation graph for subject aa.

**Figure S7:**
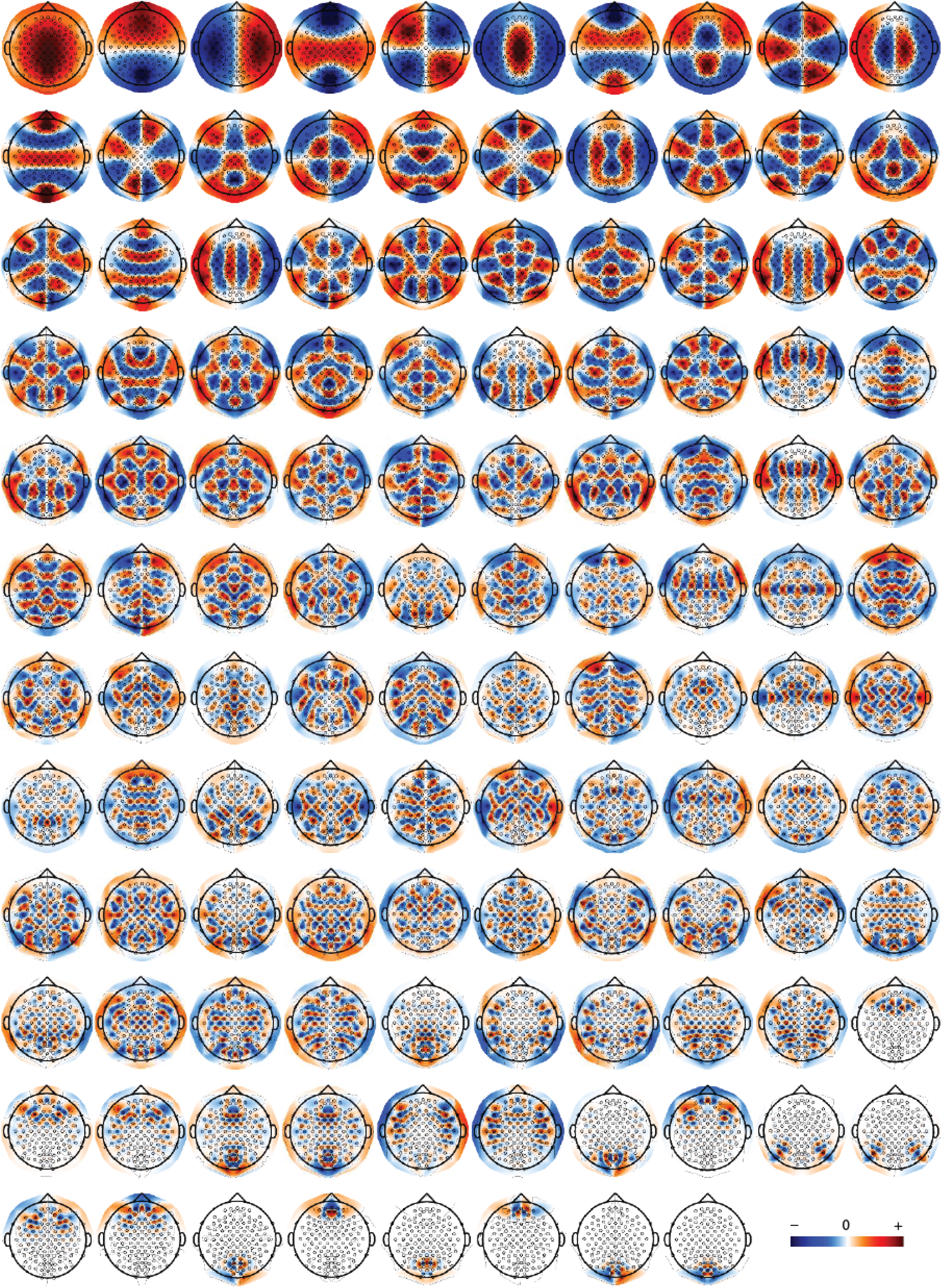
The Laplacian eigenmaps of the Euclidean distance graph for subject aa.

**Figure S8:**
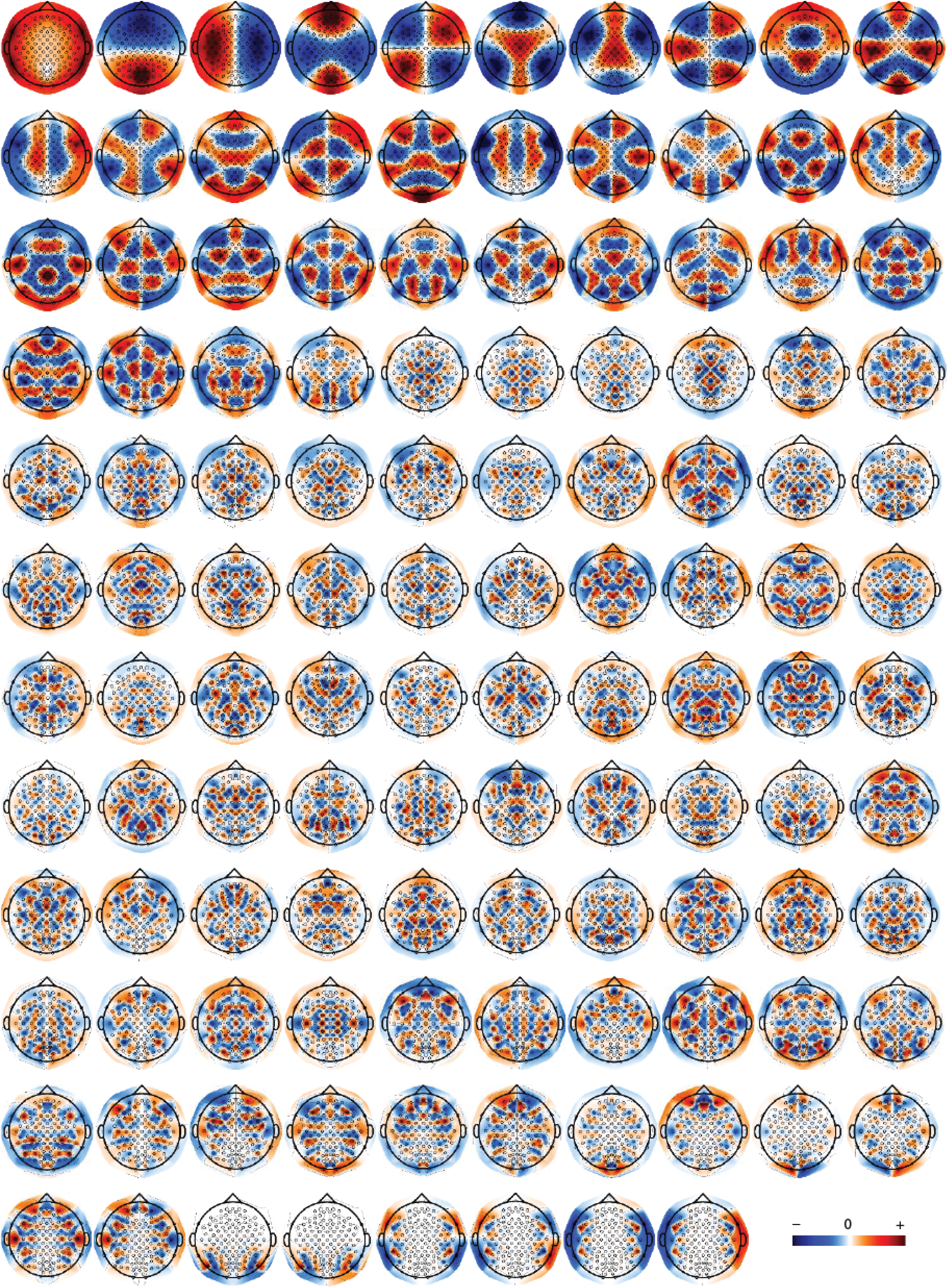
The Laplacian eigenmaps of the Euclidean distance-thresholded graph for subject aa.

**Figure S9:**
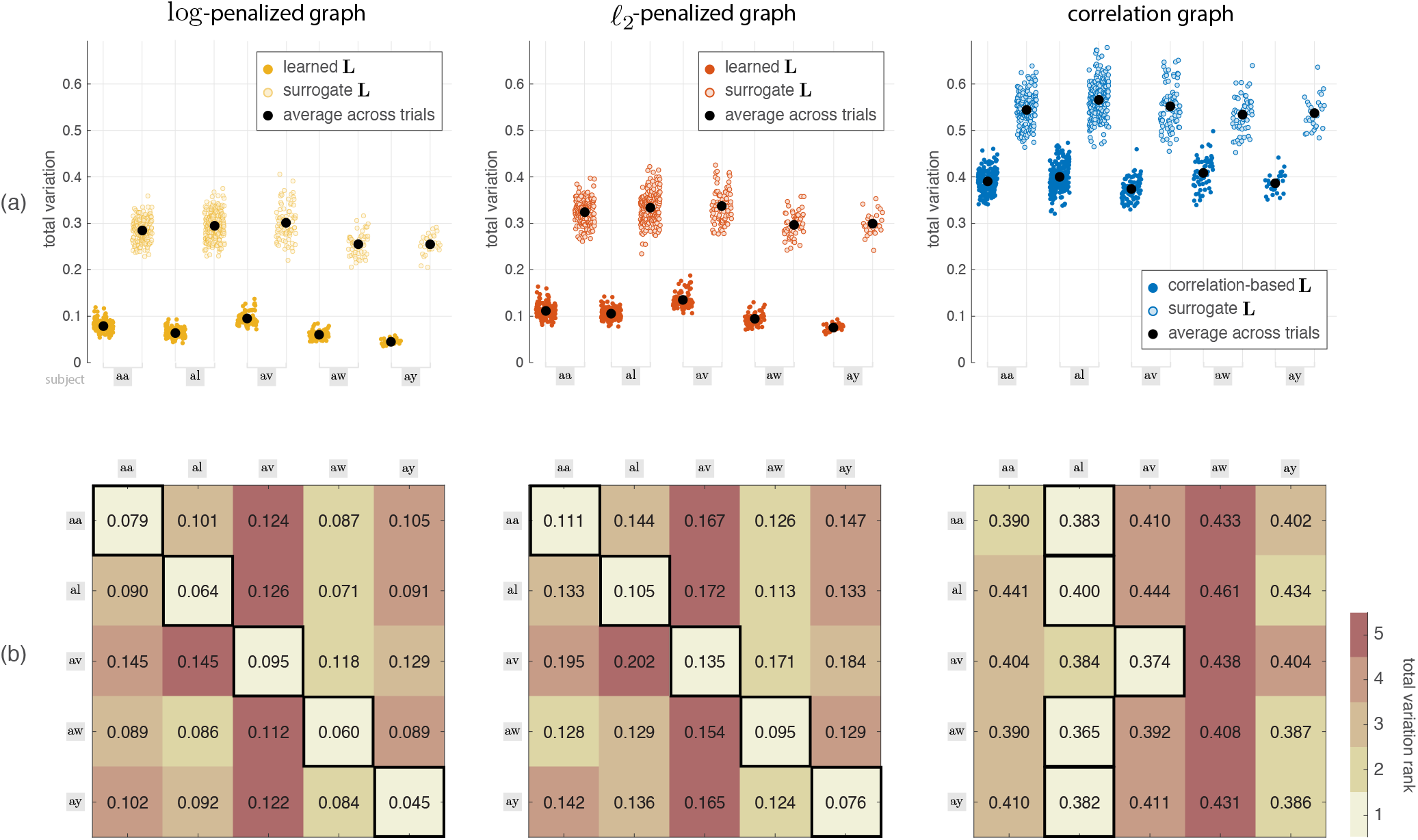
Same as in Figure 6 but using the training-set data, i.e., data that was used to learn the graphs. For the learned graphs, values are lower than those in Figure 6, as expected, since the graphs are more adapted to the data that is being tested. For the same reason, for the correlation graph, the mean TV values of the first four subjects are lower than those obtained on the test-set data, whereas the values are higher in subject ay that may be related to the small number of training-set data in this subject compared to the other subjects.

**Figure S10:**
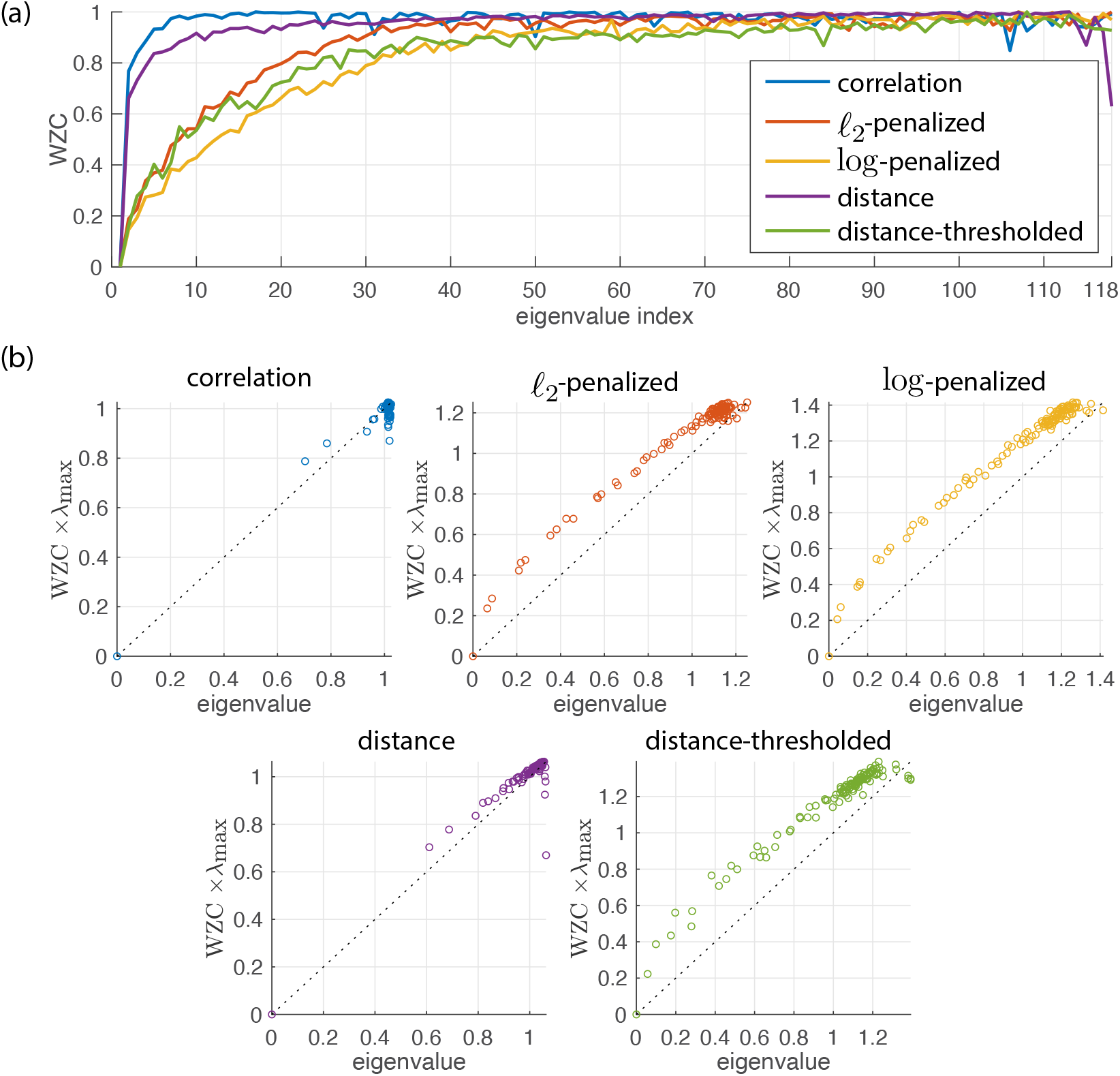
(a) Weighted zero crossing measure for the eigenvectors of five different graphs for subject aa. (b) Relation between the WZC measure of the eigenvectors and their corresponding eigenvalues. In the distance graph which is a complete graph, WZC shows a behavior similar to the correlation graph, and the WZC of the distance-thresholded graph which is a much sparser graph, has a trend similar to the learned graphs.

**Figure S11:**
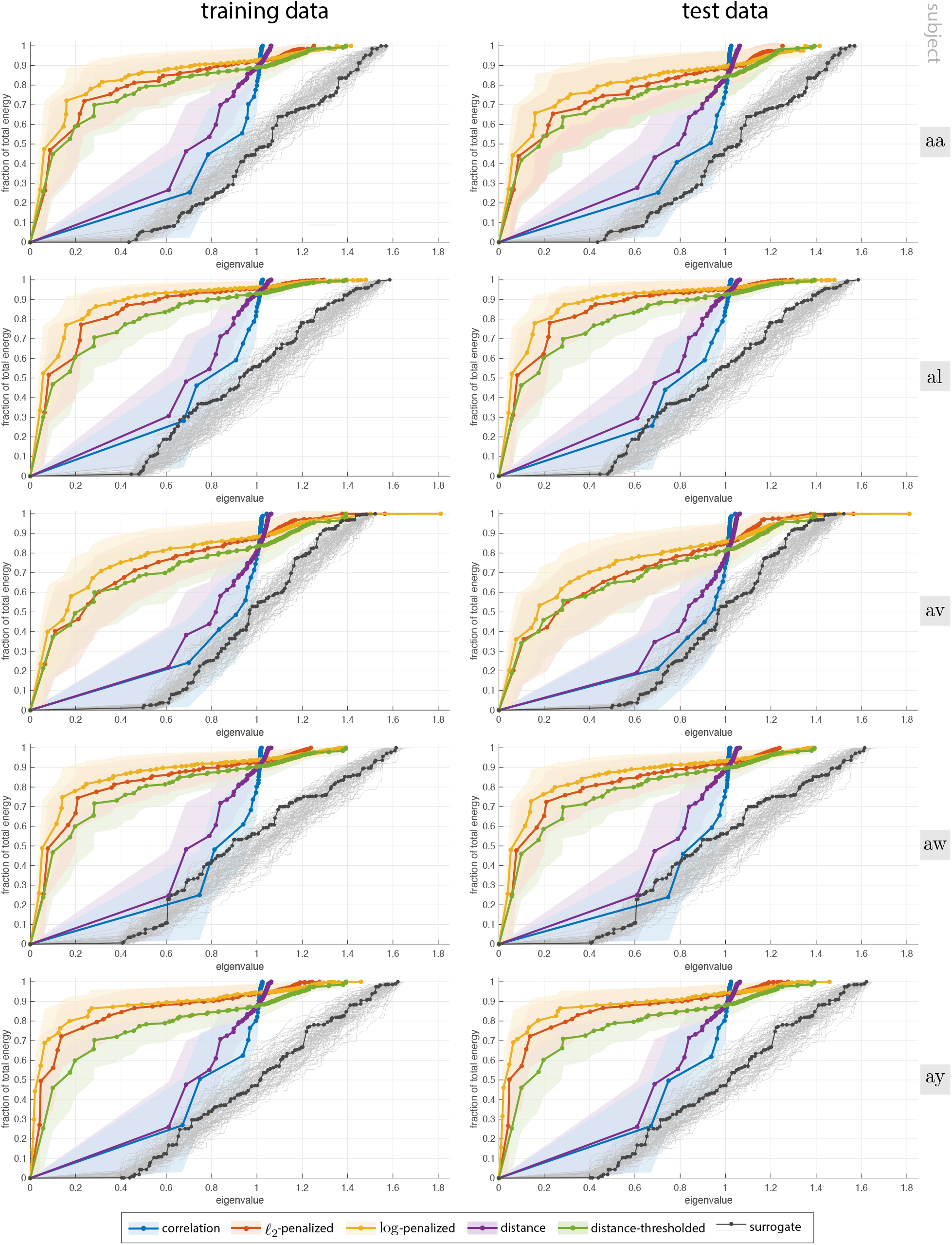
Cumulative energy spectra of EEG graph signals on five different graphs, on training- and test-set data, across five subjects. The two distance graphs are identical across subjects. For each subject, the three functional graphs (correlation and two learned graphs) are build using the training data, using the Laplacian harmonics of which the cumulative energy spectra are computed for both the training- and test-data sets. Results are also presented on 100 surrogate graphs, generated using the same scheme as described for Figure 8. Given that each surrogate graph has a unique spectrum, results are presented as individual curves rather being averaged; the eigenvalues of a representative surrogate graph are marked.

**Figure S12:**
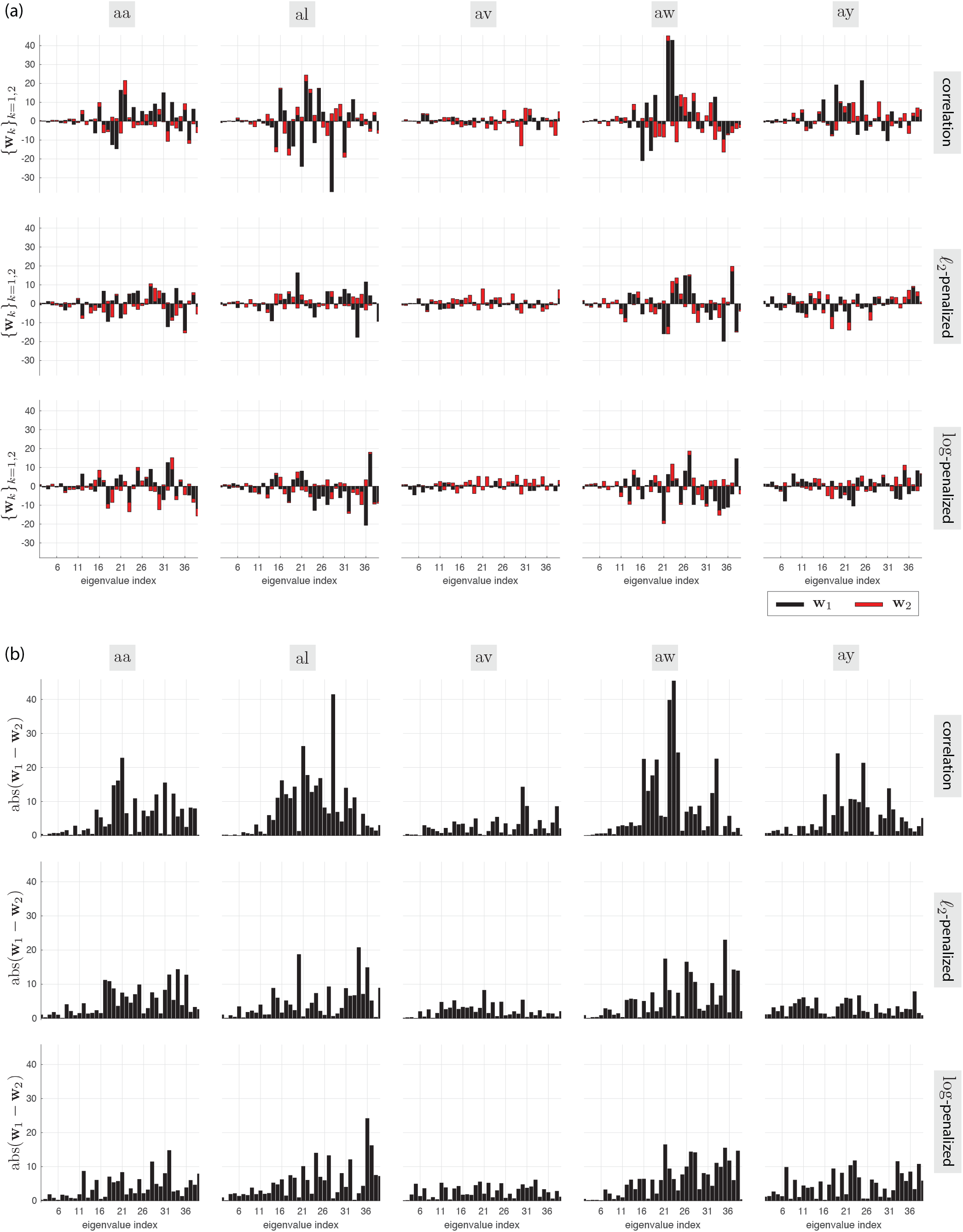
(a) FKT filters for the three studied graphs of all five subjects using the lower end of their spectra. (b) The absolute value difference between the two filters in each subject for each graph, indicating the harmonics which provide maximal contribution for discrimination the two classes.

**Table S1:**
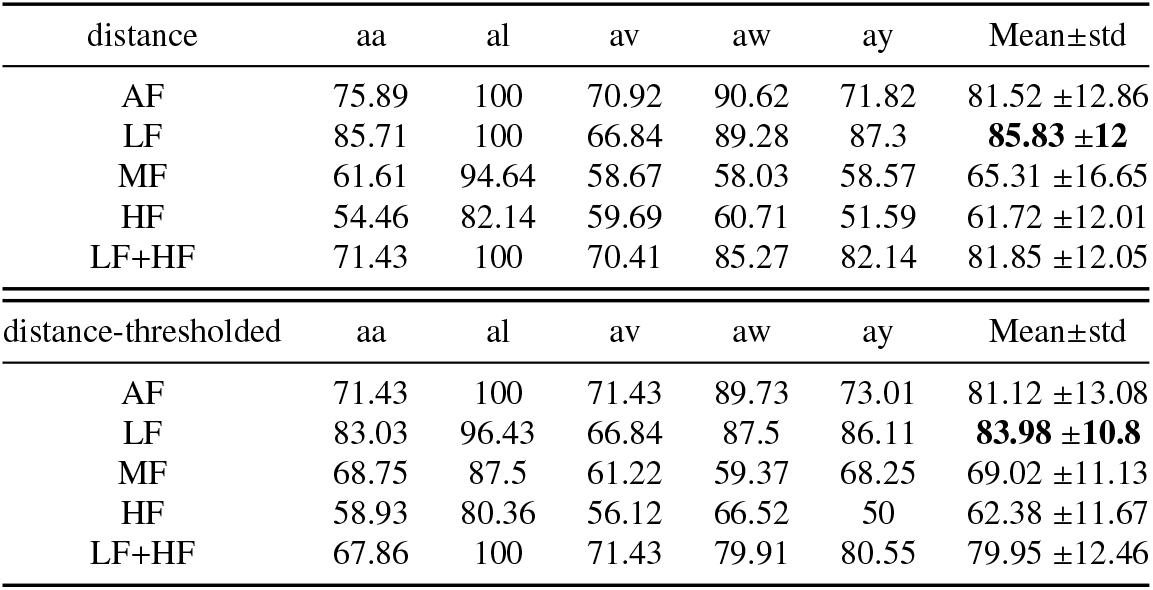
Classification accuracies (in %) on the test-set data for each subject and on average across subjects using Euclidean brain graphs, in five different frequency band settings.

| distance | aa | al | av | aw | ay | Mean±std |
| --- | --- | --- | --- | --- | --- | --- |
| AF | 75.89 | 100 | 70.92 | 90.62 | 71.82 | 81.52 ±12.86 |
| LF | 85.71 | 100 | 66.84 | 89.28 | 87.3 | **85.83 ±12** |
| MF | 61.61 | 94.64 | 58.67 | 58.03 | 58.57 | 65.31 ±16.65 |
| HF | 54.46 | 82.14 | 59.69 | 60.71 | 51.59 | 61.72 ±12.01 |
| LF+HF | 71.43 | 100 | 70.41 | 85.27 | 82.14 | 81.85 ±12.05 |
| distance-thresholded | aa | al | av | aw | ay | Mean±std |
| AF | 71.43 | 100 | 71.43 | 89.73 | 73.01 | 81.12 ±13.08 |
| LF | 83.03 | 96.43 | 66.84 | 87.5 | 86.11 | **83.98 ±10.8** |
| MF | 68.75 | 87.5 | 61.22 | 59.37 | 68.25 | 69.02 ±11.13 |
| HF | 58.93 | 80.36 | 56.12 | 66.52 | 50 | 62.38 ±11.67 |
| LF+HF | 67.86 | 100 | 71.43 | 79.91 | 80.55 | 79.95 ±12.46 |

**Table S2:** Classification accuracies (in %) using the distance graphs when directly using the GFT coefficients vs the proposed method. In both settings, features are selected from the LF sub-band of the graph spectra.

| GFT | aa | al | av | aw | ay | Mean±std |
| --- | --- | --- | --- | --- | --- | --- |
| distance | 66.96 | 92.86 | 55.61 | 64.73 | 63.49 | 68.73 ±14.15 |
| distance-thresholded | 66.07 | 80.36 | 66.84 | 66.07 | 60.71 | 68.01 ±7.33 |
| Proposed | aa | al | av | aw | ay | Mean±std |
| distance | 85.71 | 100 | 68.88 | 89.28 | 87.7 | 86.31 ±11.21 |
| distance-thresholded | 83.93 | 100 | 67.86 | 87.5 | 94.84 | **86.82 ±12.31** |

## Footnotes

1 Further details about this dataset can be found at http://www.bbci.de/competition/iii/)

